# Rapid Detection of Genetic Engineering, Structural Variation, and Antimicrobial Resistance Markers in Bacterial Biothreat Pathogens by Nanopore Sequencing

**DOI:** 10.1101/730093

**Authors:** A. S. Gargis, B. Cherney, A. B. Conley, H. P. McLaughlin, D. Sue

## Abstract

Widespread release of *Bacillus anthracis* (anthrax) or *Yersinia pestis* (plague) would prompt a public health emergency. During an exposure event, high-quality whole genome sequencing (WGS) can identify genetic engineering, including the introduction of antimicrobial resistance (AMR) genes. Here, we developed rapid WGS laboratory and bioinformatics workflows using a long-read nanopore sequencer (MinION) for *Y. pestis* (6.5h) and *B. anthracis* (8.5h) and sequenced strains with different AMR profiles. Both salt-precipitation and silica-membrane extracted DNA were suitable for MinION WGS using both rapid and field library preparation methods. In replicate experiments, nanopore quality metrics were defined for genome assembly and mutation analysis. AMR markers were correctly detected and >99% coverage of chromosomes and plasmids was achieved using 100,000 raw sequencing reads. While chromosomes and large and small plasmids were accurately assembled, including novel multimeric forms of the *Y. pestis* virulence plasmid, pPCP1, MinION reads were error-prone, particularly in homopolymer regions. MinION sequencing holds promise as a practical, front-line strategy for on-site pathogen characterization to speed the public health response during a biothreat emergency.

## Introduction

*Yersinia pestis* and *Bacillus anthracis* are the etiological agents of plague and anthrax, respectively. Deliberate misuse of these pathogens as bioterrorism agents poses a serious public health and safety threat, due to low infectious doses, high lethality, and the ease of production, dissemination and communicability^1^. Strains of both species were weaponized previously, and the 2001 Amerithrax incident underscores the importance of biological threat preparedness and rapid laboratory response^2, 3^. Natural anthrax and plague outbreaks are reported annually in animals, and human cases are often zoonotic. While these diseases continue to cycle in isolated, enzootic foci, re-emergence was reported during 2016 anthrax outbreaks in Russia and Sweden^4^, and the 2017 pneumonic plague outbreak in Madagascar^5^. Both naturally-occurring and engineered antimicrobial resistance (AMR) have been reported in *Y. pestis* and *B. anthracis*^6^. Most *B. anthracis* strains are susceptible to antibiotics, but surveys of clinical and environmental isolates indicate penicillin resistance occurs in 2 to 16% of strains^7^ Attenutated *B. anthracis* strains with AMR were isolated following laboratory *in vitro* passaging (selective pressure) and/or targeted genetic manipulation^8, 9^ Laboratory-selected virulent and avirulent *Y. pestis* strains with quinolone-resistance were previously reported^10, 11^ and engineered multi-drug resistant (MDR) strains are a public health concern^12, 13^. *Y. pestis* is intrinsically susceptible to every antibiotic recommended for human plague therapy, but a few *Y. pestis* isolates with transferable plasmid-mediated resistance were reported from Madagascar^13, 14, 15, 16^.

A public health response to incidents involving human plague or anthrax requires timely and accurate AMR profiling. Conventional broth microdilution (BMD) is the gold standard antimicrobial susceptibility testing (AST) method, but requires lengthy incubation times for *B. anthracis* (16 to 20 hours) and *Y. pestis* (at least 24 hours)^17, 18^. Several new phenotypic methods can rapidly assess functional susceptibility using: real-time PCR to detect growth in the presence of antimicrobial agents^19^, bioluminescent reporter phage^20^, flow cytometry^21^, laser light scattering technology^22^, and optical screening^23^. While these phenotypic methods are more rapid than BMD AST, they cannot definitively detect genetic manipulation. Whole genome sequencing (WGS) can reveal evidence of genetic engineering in bacteria, such as the introduction of genes, mutations and/or plasmids. While there is not a comprehensive understanding of all genes or mutations associated with resistance, WGS is invaluable for understanding novel genotypic/phenotypic relationships. Moreover, unlike PCR–based methods, a comprehensive knowledge of AMR is not required, since *de novo* WGS is unbiased and can reveal previously undescribed markers without targeted primer sets. Genomic data complements phenotypic AST results and offers insight into mechanisms of drug resistance, which is especially critical in the age of synthetic biology.

Two next-generation sequencing (NGS) approaches are used for microbial WGS: short-read sequencing (e.g. Illumina) and long-read-sequencing (e.g. Pacific Biosciences, PacBio and Oxford Nanopore Technologies, ONT). Short-read sequence data have low error rates and are useful for strain typing and the identification of single nucleotide polymorphisms (SNPs)^24, 25^. The short (maximally 300nt) read lengths limit the ability to resolve repetitive sequences or structural genomic rearrangements, and can result in unfinished, fragmented *de novo* assemblies comprised of hundreds of contigs^25, 26^. Many AMR gene regions are flanked by repetitive insertion sequences (IS), and short-read sequencing cannot span these regions^25^. This can result in inaccurate chromosome or plasmid assemblies, including multiple copies of a repeat region collapsed into one location^25, 27^ Long-read sequencing platforms produce reads ranging from several kilobases (kb) to greater than 10 kb in length that can bridge gaps between repetitive regions and allow complete bacterial genome finishing^25^. However, long-read sequence data have higher error rates than short-read data^28^, and are dominated by insertions and deletions (indels), especially in homopolymeric regions^24, 29^ Small plasmids (<7 kb) are difficult to accurately assemble from PacBio reads due to size selection limitations of library preparation and data analysis^30, 31^. However, short- and long-read NGS data are complementary and together generate highly accurate hybrid genome assemblies^25, 26, 32, 33^.

Work with *Y. pestis* and *B. anthracis* requires specialized facilities to ensure both biosecurity and biosafety. Physical, high-containment laboratory space is often limited, and instruments used for molecular characterization, such as WGS, are typically located in separate areas at lower containment levels. NGS instruments can incur high capital costs, occupy a large footprint or require specialized technical training, making routine on-site sequencing cost- and space-prohibitive^28, 34^. By contrast, there is minimal capital cost for the MinION (ONT) device, which is the size of a USB thumb-drive, and its operation requires a computer or ONT device controller that together occupy little benchtop space. Therefore, nanopore WGS can occur within a high-containment laboratory with limited space. Unlike PacBio sequencing that requires ≥1 μg of genomic DNA (gDNA), only 400 ng of gDNA is required for library preparation using the MinION Rapid Sequencing Kit. Sequencing libraries can also be prepared using a MinION Field Sequencing Kit that includes shelf-stable lyophilized reagents. After a run begins, MinION sequence data are available in minutes enabling real-time sequence analysis.

The field of nanopore sequencing is rapidly evolving, and benchmarking studies that use standardized conditions to assess run-to-run variability are lacking. Few studies have assessed the accuracy and consistency of nanopore sequencing using the same strain with consistent chemistry, consumables, and software versions^26, 35, 36^. We developed a custom bioinformatics pipeline to compare the genome assemblies from standardized replicate nanopore sequencing experiments to define sequencing quality metrics, establish performance characteristics, and demonstrate the feasibility of rapid long-read WGS for *Y. pestis* and *B. anthracis*. The resulting same-day laboratory workflow for DNA isolation, library preparation, nanopore sequencing, and bioinformatics analysis is timely and effective for the rapid detection of known markers related to AMR and evidence of genetic engineering in bacterial biothreat pathogens.

## Results

### *Y. pestis* gDNA quality and quantity

All *Y. pestis* gDNA met ONT criteria for MinION sequencing as measured by optical density absorbancy ratios at 260 nm and 280 nm (A260/280) and at 260 nm and 230 nm (A260/230) (Supp. Table 1). Both silica-membrane (SM)- and salt-precipitation (SP)-based gDNA extraction methods produced *Y. pestis* gDNA of comparable quality and quantity, with intact high molecular weight (HMW) and little evidence of shearing (Supp. Fig. 1). The SP method produced higher molecular weight DNA compared to SM extraction, but required an hour longer to complete.

### *Y. pestis* MinION sequencing technical replicates using Rapid Sequencing and Field Sequencing Kit libraries

*Y. pestis* S1037, a derivative of the attenuated strain All22^37^, was sequenced to establish baseline performance and quality metrics for rapid nanopore WGS. Strain S1037 contains two virulence plasmids, pMT1 (~110 kb) and pPCP1 (9.5 kb monomer and ~19 kb dimer), and is ciprofloxacin non-susceptible (CIP-NS, MIC 4 μg/ml by BMD AST). Unlike A1122, strain S1037 contains a *gyrA* single nucleotide polymorphism (SNP) (Table 1) in the previously described *Y. pestis* quinolone resistance-determining region (QRDR)^11^. Three replicate rapid sequencing libraries were prepared from one SP gDNA extraction (S1037, SP, Rapid 1, 2, and 3; Fig. 1) and from one SM gDNA extraction (S1027, SM, Rapid 1, 2, and 3; Fig. 1) to evaluate variation among sequencing runs from the same sample. Libraries were also prepared with the Field Sequencing Kit using the same SP and SM gDNA used in the technical replicate study (SP Field and SM Field; Fig. 1). Each library was sequenced for up to 48 hours, but only reads generated during the first 16 hours of sequencing were analyzed to allow for a standardized comparison of data sets.

**Figure 1.**
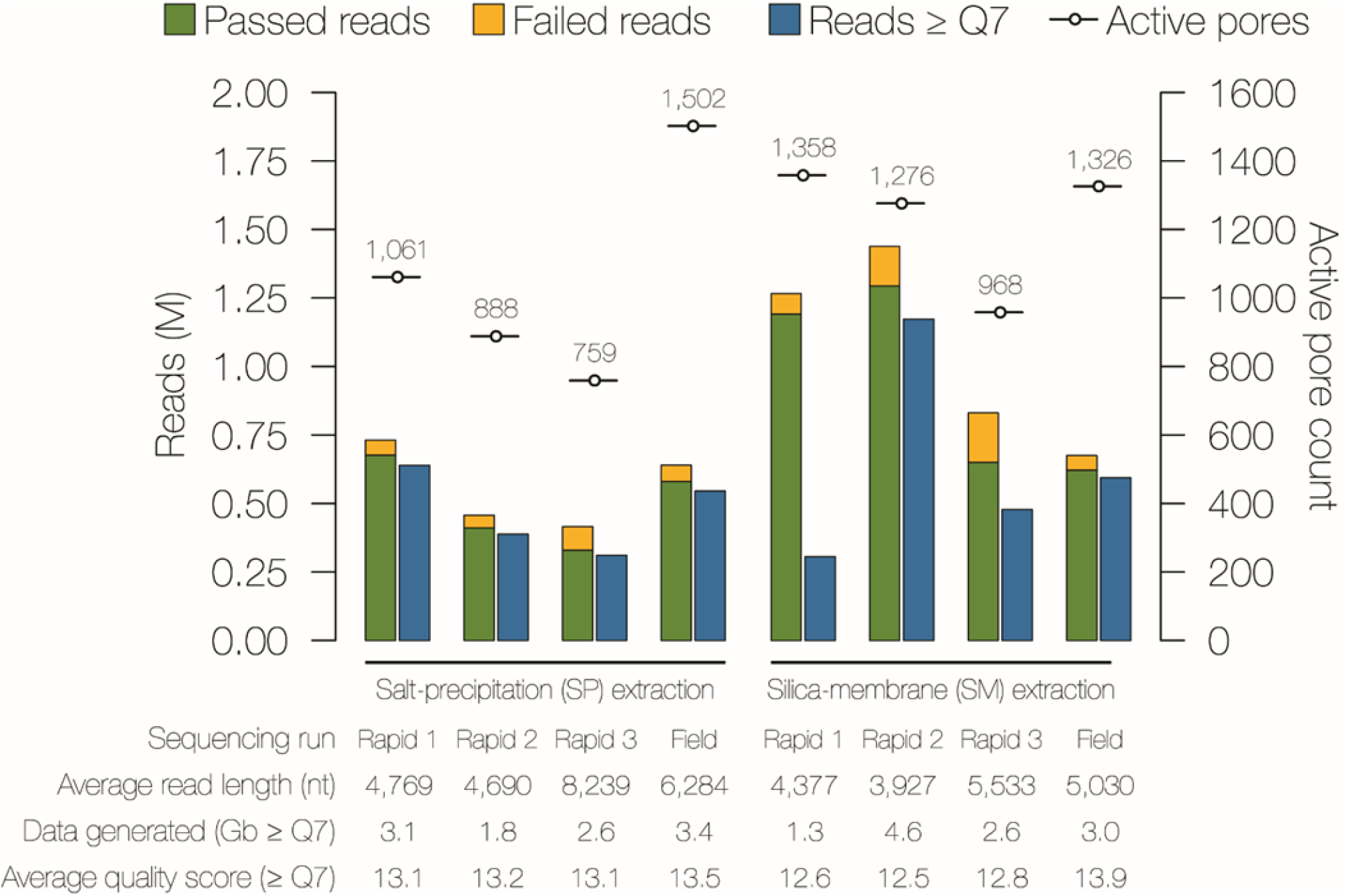
Nanopore sequencing run metrics for *Y. pestis* strain S1037. Genomic DNA was extracted by the salt-precipitation (SP) or the silica-membrane (SM) method and libraries were prepared in triplicate using the ONT Rapid Sequencing Kit (Rapid), or the Field Sequencing Kit (Field). All run metrics were based on a 16 hour sequencing run time and the active pore count (-o-) was recorded during the platform QC step prior to the sequencing run. The number of raw fast5 reads generated (green, passed; yellow, failed) and raw FASTQ Albacore base-called reads with an average quality score of ≤ 7 (blue) are displayed.

**Table 1.**
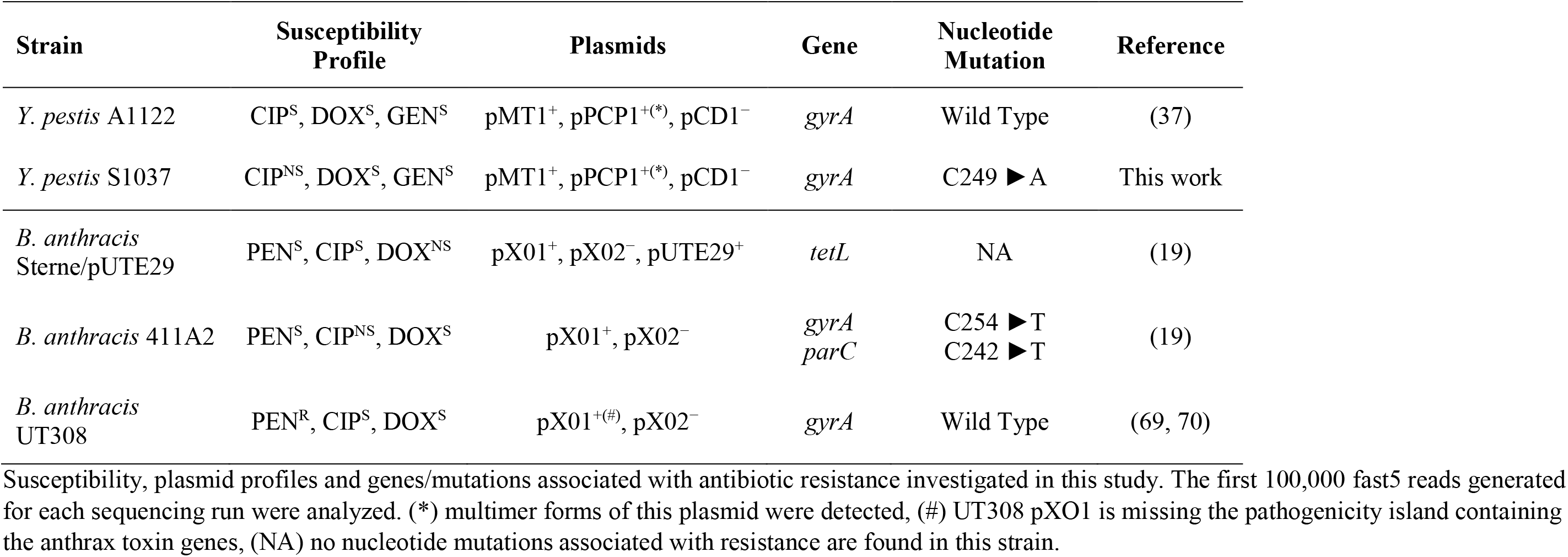
Bacterial strains used in this study.

After base calling, FASTQ reads with an average quality (Q) score of <7 were discarded, resulting in removal of 4 to 74% of total reads from analysis (Fig. 1, Supp. Table 3). The MinKNOW software became unresponsive after ~11-12 hours of sequencing for SP Rapid 1 and SM Rapid 1, and both were restarted and continued for ~6-8 additional hours (Supp. Table 2). While SP Rapid 1 appeared unaffected by the software problem, SM Rapid 1 lost ~74% of reads following basecalling and contained the fewest reads ≥Q7 of any data set. All flowcells met ONT’s minimum active pore requirement (≤800) during platform QC, but the flowcell used for SP Rapid 3 had an initial QC of 871 active pores that decreased to 759 after loading the library (Fig. 1, Supp. Table 3). SP Rapid 3 lost ~6% of reads following base-calling, but the amount of data generated and average Q-score resembled other technical replicate runs (Fig. 1, Supp. Table 3).

The number of ≥Q7 FASTQ reads generated was inconsistent within and among the SP and SM replicate runs. During the first 16 hours of sequencing, the SP technical replicates generated 310,906-639,598 >Q7 reads, while SM experiments generated 305,824-1,172,803 ≥Q7 reads. The average read lengths within and among the SM and SP replicates varied. Both average read length and number of reads generated contributed to the amount of useable data (Fig. 1, Supp. Table 3). For example, SP Rapid 2 (388,435 reads) had more ≥Q7 reads than SP Rapid 3 (310,906 reads), but SP Rapid 3 read lengths were ~1.75X longer. The amount of SP Rapid 3 run data generated was greater (2.6 Gb vs. 1.8 Gb). Overall, the range of useable data generated by the SP replicate experiments (1.8 to 3.1 Gb) was comparable to SM (1.3 to 4.6 Gb). The average Q-scores of the SM and SP Rapid Sequencing technical replicates ranged from 12.5 to 13.2. The amount of sequencing data (≥Q7 reads and total Gb) generated using the Field Sequencing Kit was comparable to the Rapid Sequencing Kit (Fig. 1, Supp. Table 3). The Field Sequencing Kits produced data sets with the overall highest SP (13.5) and SM (13.9) Q-scores.

The strand-to-pore ratio is a quality metric that represents the quantity of MinION flow cell pores actively sequencing DNA (pores in strand) relative to total available pores. The ratio was recorded after ~10 minutes of sequencing per run to assess the library preparation quality (Supp. Table 2). A ≥50% strand-to-pore ratio was expected for the RAD-004 Rapid Sequencing Kit (ONT Community and personal communication). For SP Rapid 1, 2, and 3, the average strand-to-pore ratios were 60%, 63%, and 64%, respectively; and for SM Rapid 1, 2, and 3, the ratios were 80%, 75%, and 82%, respectively. Ratios increased to >80% within the first 1-2 hours of sequencing for all SM and SP replicates.

### MinION sequencing performance and *Y. pestis* S1037 *de novo* assembly using retrospective downsampling of passed fast5 reads

A retrospective downsampling of passed fast5 reads was performed to determine the amount of data necessary to produce high-quality genome assemblies. Assembly quality was measured by indel frequency, genome coverage, ability to assemble large and small plasmids, and accurate detection of the *gyrA* SNP associated with the CIP-NS phenotype (Fig. 2).

**Figure 2.**
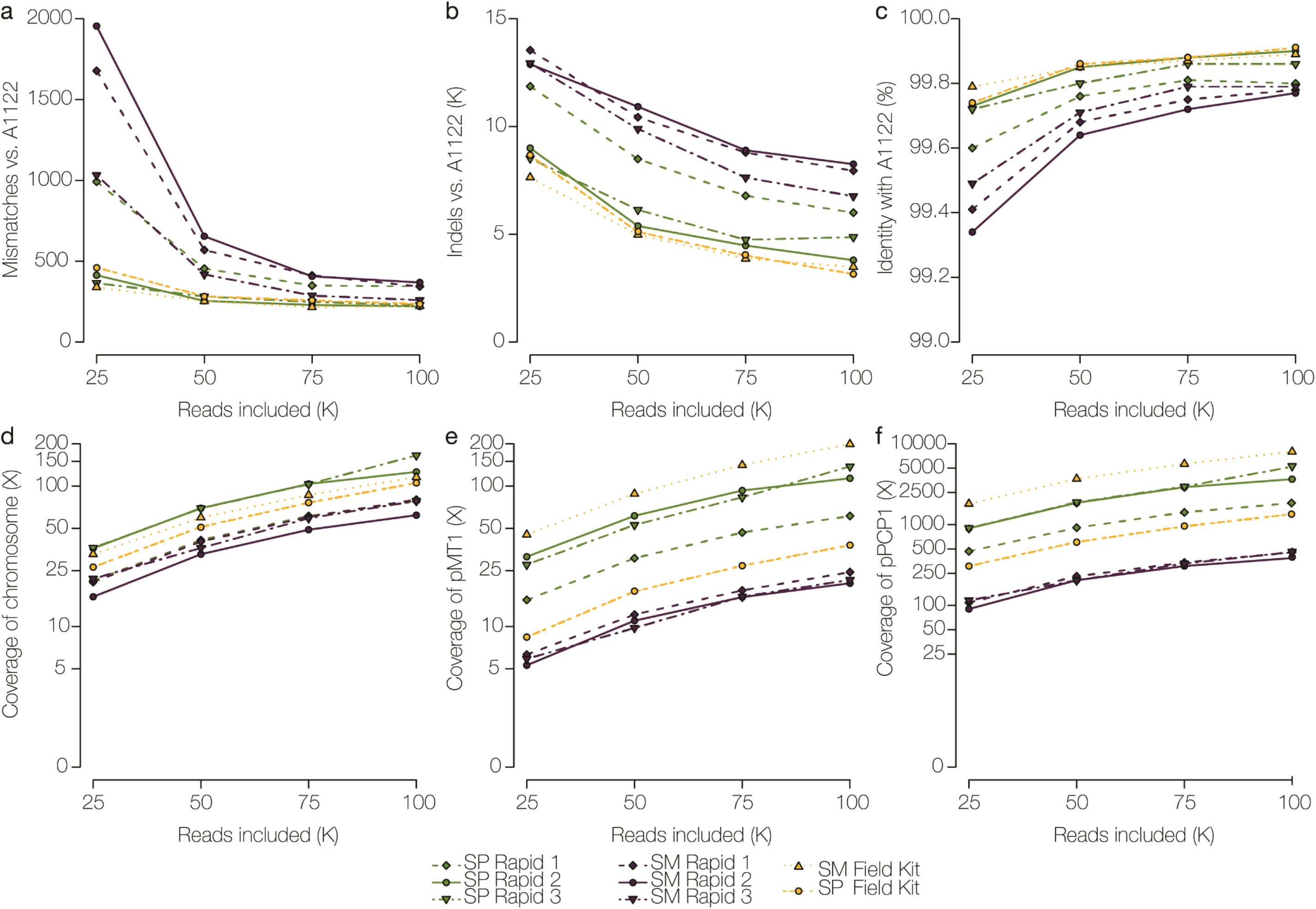
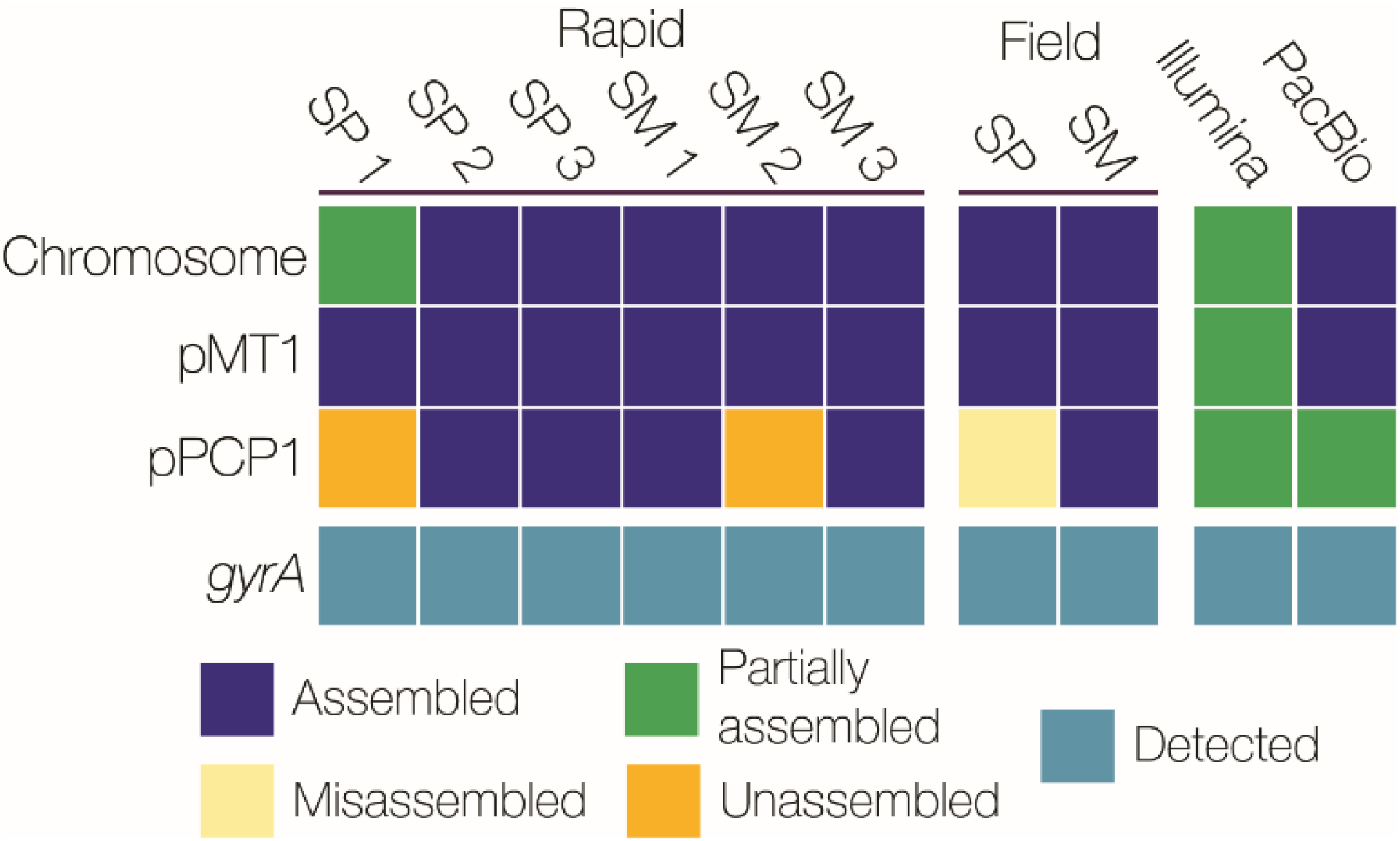
Assessment of nanopore sequencing performance and *de novo* assembly for *Y. pestis* S1037. DNA sequencing libraries were prepared using the Rapid Sequencing Kit (in triplicate) and Field Sequencing Kit from a salt-precipitation (SP) or a silica-membrane (SM) extraction. Assemblies were generated using analysis of the first 25,000, 50,000, 75,000, and 100,000 passed fast5 nanopore reads. **a-c** Determination of mismatches, indels, and % identity, and **d-f** fold coverage for chromosomal and plasmid sequences of *Y. pestis* S1037 data sets aligned to the All22 reference strain. **g** Using 100,000 passed fast5 reads, the ability to assemble the chromosome and plasmids, and to detect the *gyrA* SNP associated with the non-susceptible CIP phenotype is depicted in a presence/absence plot. Assembled is defined as a complete sequence with the same nucleotide order as the reference sequence (navy), partially assembled represents a sequence that was assembled but not as a single contig (green), misassembled includes a sequence that was assembled but the bases are in a different order than the reference sequence, or a sequence assembled multiple times resulting in an incomplete assembly (yellow), and unassembled refers to a sequence that was not present in the assembly (orange).

Minimal improvements in overall assembly quality were observed when >75,000 reads were analyzed (Fig. 2). *De novo* assemblies using ≤75,000 reads did not consistently assemble the *Y. pestis* chromosome, or large (pMT1) and small (pPCP1) plasmids. Subsequent nanopore analyses were performed with 100,000 reads only. The average number of mismatches for the SP and SM 100,000-read data sets ranged from 220 to 368, while Illumina MiSeq and PacBio assemblies of the same strain contained 44 and 29 mismatches, respectively (Fig. 2a). Thousands of indels (3,151 to 8,251) were detected in nanopore assemblies (Fig. 2b), compared to Illumina (18 indels) and PacBio (93 indels), and were predominantly located in homopolymeric regions. Indels were not significantly reduced in nanopore assemblies supplemented with >100,000 additional reads (data not shown). Using 100,000 reads, averaged across the three Rapid SP and three Rapid SM *Y. pestis* S1037 data sets, the poly-N length and the fraction of poly-N’s correct in the assemblies was plotted for each nucleotide type using both the RACON and Nanopolish error corrected assemblies (Supp. Fig. 2). Nanopolish increased the fraction of *Y. pestis* poly-N lengths correctly called, but this fraction exponentially decreased for lengths >4. Increased accuracy was observed when calling poly-As and Ts compared to poly-Gs and Cs. The *gyrA* point mutation, which is not located in a homopolymeric region, was correctly identified in all *Y. pestis* assemblies (Fig. 2g).

The first 100,000 fast5 reads yielded >99% identity with A1122 across the ~4.6 megabase *Y. pestis* genome for all experiments (Fig. 2c). For the chromosome, all nanopore assemblies generated a single contig, except the SP Rapid 1 data set (2 contigs observed) (Fig. 2g). The average depth of chromosome coverage was 101X for all replicate *Y. pestis* S1037 data sets (Fig. 2d). Plasmid pMT1 was consistently assembled to completion, with multiple overlaping reads spanning the entire plasmid length in all 100,000-read data sets, however pPCP1 was not. Un-or misassembled pPCP1 fragments were detected (i.e. assembly of multiple small fragments with homology to pPCP1 or bases in different order than the reference sequence) (Fig. 2g). Reads corresponding to pPCP1 with lengths greater than 9.5 kb and 19 kb were also identified, and are discussed below. Overall, the SP sequencing runs produced higher average coverage for both pMT1 (104X) and pPCP1 (3,567X) compared to the SM runs (22X and 431X, respectively) (Fig. 2e and 2f). Among the SM and SP Rapid and Field Sequencing Kit runs, the average time to generate 100,000 fast5 reads was ~2 hours (Supp. Table 2) and data analysis (including base-calling) required ~2.5 additional hours. For *Y. pestis*, SM gDNA extraction from an agar culture isolate, MinION WGS, *de novo* assembly and analysis can be completed in ≤6.5 hours (Fig. 3).

**Figure 3.**
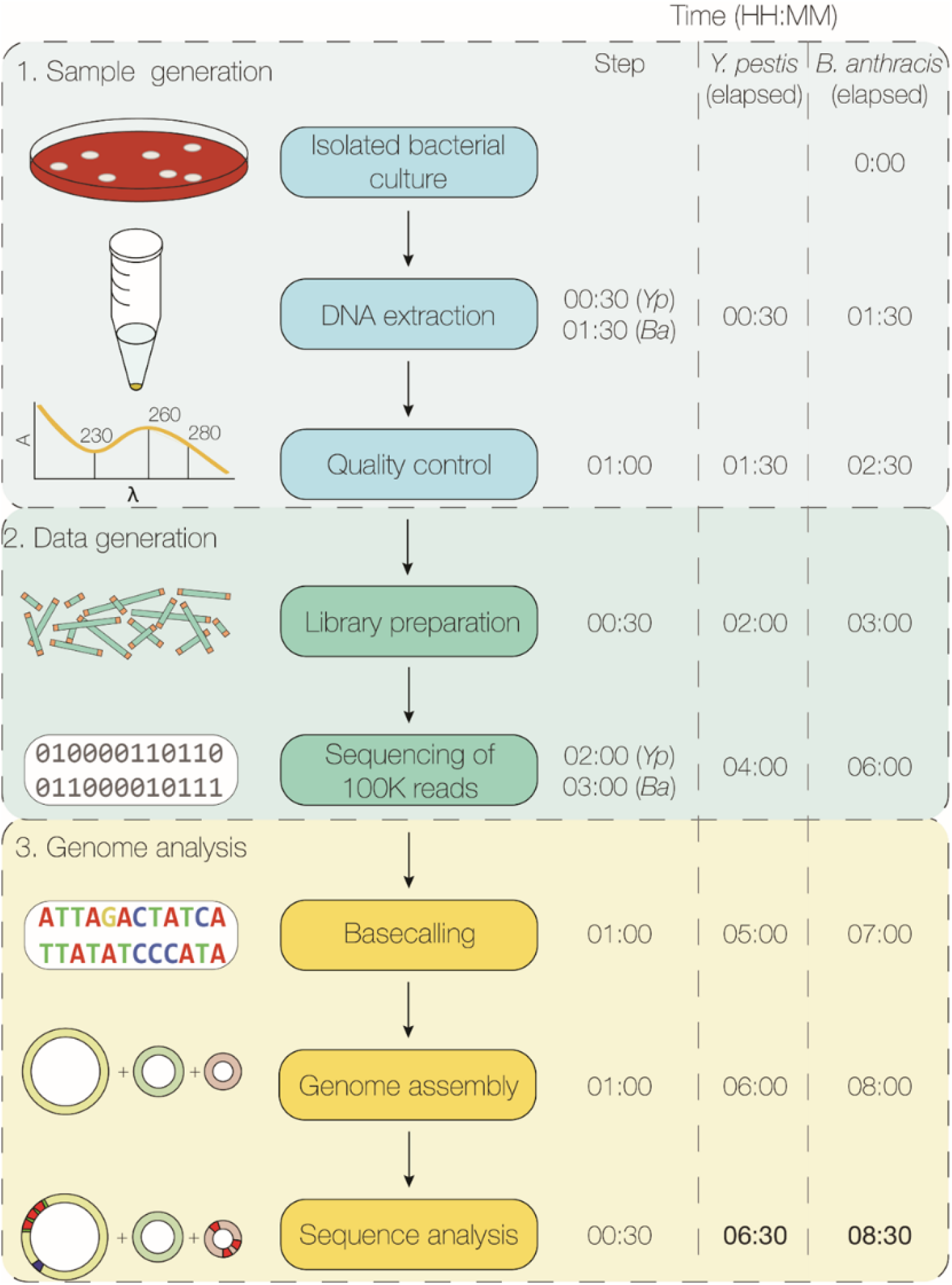
Time required to detect antimicrobial resistance markers in *Y. pestis* and *B. anthracis* using nanopore whole genome sequencing. Time 0:00 starts from a pure isolated culture. The time to extract genomic DNA by the silica-membrane method is shown, including the one hour *B. anthracis* lysis step. Quality control steps include assessment of DNA concentration, quality, and integrity. Library preparation time is estimated for the Rapid Sequencing Kit. The average time required to generate 100,000 passed fast5 reads is shown for each species. Genome analysis includes base-calling using Albacore and *de novo* genome and plasmid assembly. See methods and Supplementary Figure 9 for the bioinformatics analysis pipeline details.

### Detection of multimeric pPCP1 forms in *Y. pestis* S1037 and A1122 assemblies

Analysis of read length distributions among the SP and SM technical replicates revealed a greater number of long reads generated in SP experiments (Supp. Fig. 3). SP data sets also contained more ~18.2 kb reads, which correspond to the pPCP1 dimer (2-mer) form. Most reads from *Y. pestis* S1037 SP data sets with regions homologous to pPCP1 were ~9.1 kb (~17%) or ~18.2 kb (~80%) (Supp. Fig. 4), which correspond to the previously described monomer (~9.5 kb) and dimer (~19 kb) forms^38^. A few reads (~3%) with lengths >18.2 kb were detected in the 100,000-read data sets from SP extracted gDNA, but were absent in SM data sets (Supp. Fig. 3 and Supp. Fig. 4). Dimer, trimer, and tetramer pPCP1 forms are linked together in a 5’-3’ head-to-toe fashion (Fig. 4). Multimers were present in the nanopore data sets derived from the Field Sequencing and three Rapid Sequencing Kit preparations of the same SP extracted gDNA (SP-1). An S1037 assembly using Illumina short-read data resulted in three contigs (688 nt, 1,894 nt, and 7,375 nt) with pPCP1 homology, and did not assemble the ≥19 kb plasmid forms (Fig. 2g). An S1037 PacBio assembly resulted in 3 contigs corresponding to the chromosome (4,576,792 nt), pMT1 (95,849 nt), and a partially assembled pPCP1 (21,732 nt) (Fig. 2g).

**Figure 4.**
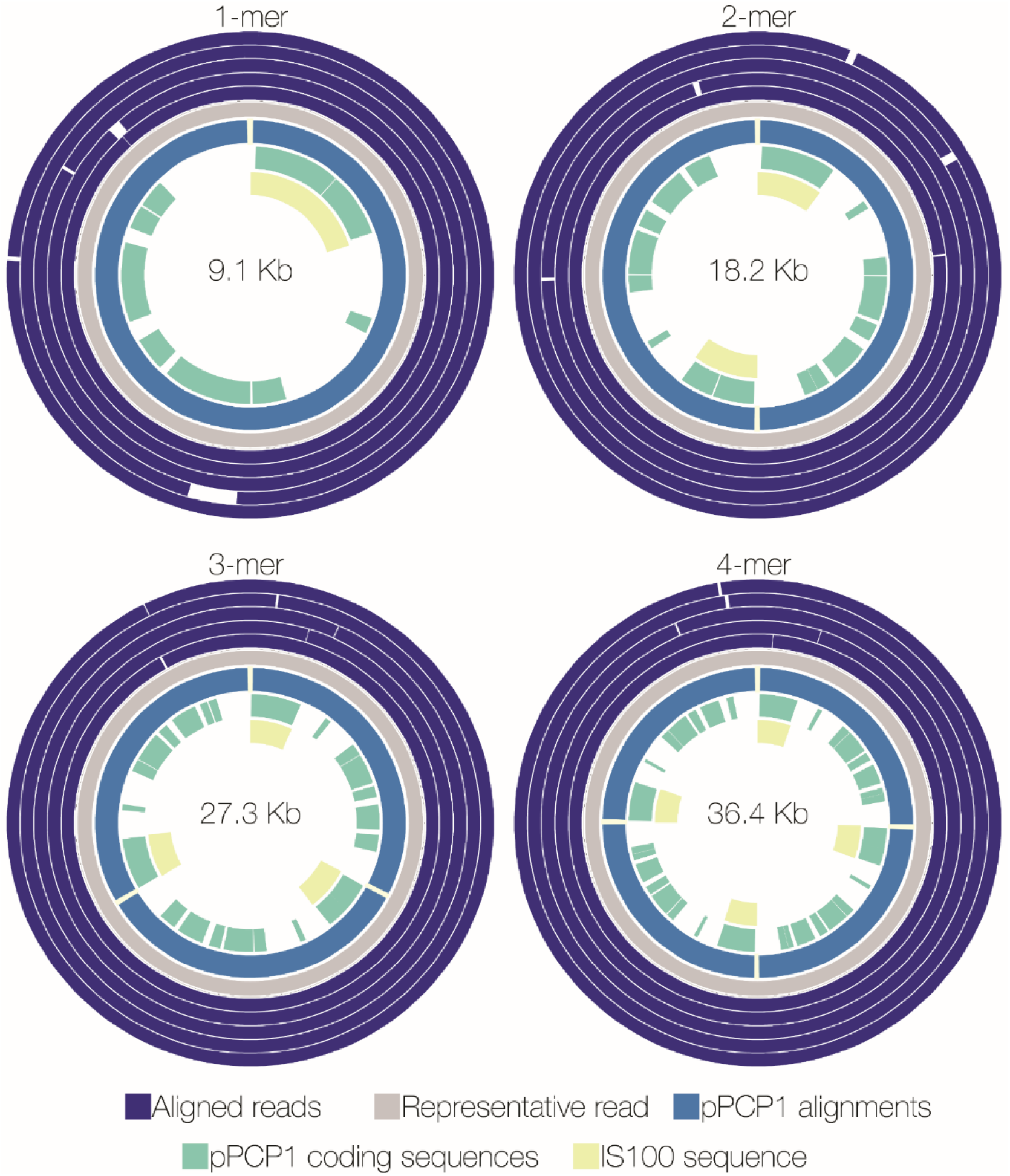
Detection of multimeric forms of pPCP1 in nanopore sequence assemblies of *Y. pestis* S1037. pPCP1 is ~9.5 kb, but the pPCP1 1-mer assembly is ~9.1 kb due to the presence of small indels in the nanopore-generated sequence. Reads >18.2 kb were assigned to the 3- or 4-mer form based on length. A representative read (grey) for each multimer was selected as a reference for the complete plasmid the remaining reads were aligned against this sequence (purple). Coding sequences (green) and IS/00 elements (yellow) are shown in the center. The 2-, 3- and 4-mer forms are linked together in a 5’-3’ head-to-toe orientation.

An additional *Y. pestis* S1037 SP gDNA extraction (SP-2, Supp. Table 1) was prepared using care to minimize DNA shearing (e.g. without vortexing and minimal pipetting) to improve WGS detection of multimer pPCP1 forms. Following sequencing using the Rapid Kit, the resulting 100,000-read d*e novo* assembly yielded comparable sequence quality to the technical replicate assemblies (data not shown) and overall contained more reads that were >18.2 kb than the S1037 SP-1 data set (Supp. Fig. 5a). The average read length (10,960 nt) was greater than the longest average read length (7,800 nt) observed in the technical replicate experiments. The SP-2 assembly contained an increased fraction of ~18.2 kb reads that corresponded to the pPCP1 plasmid dimer (Supp. Fig. 5b). Few reads with pPCP1 homology measured ~45,500 nt (4 reads) and 54,600 nt (1 read), which correspond to putative 5-mer and 6-mer forms (Supp. Fig. 5b).

To determine if *Y. pestis* A1122, the parent strain of S1037, also contained reads >18.2 kb with pPCP1 homology, SP and SM extracted gDNA was used for Rapid Sequencing Kit library preparation and nanopore sequencing. The 100,000-read *de novo* assemblies from A1122 yielded comparable sequence quality to the technical replicate assemblies (data not shown). A1122 reads >18.2 kb with pPCP1 homology (Supp. Fig 6) were assigned 3- or 4-mer forms. When >100,000 reads were analyzed, additional pPCP1 homologous reads in this size range or greater were not detected. A PacBio assembly of A1122 resulted in 3 contigs corresponding to the chromosome (4,586,979 nt), pMT1 (96,144 nt), and pPCP1 (8,935 nt) (data not shown).

### *B. anthracis* gDNA quality and quantity

The rapid *Y. pestis* nanopore sequencing approach was applied to *B. anthracis* strains with different AMR profiles to evaluate *de novo* assembly quality and to identify plasmids and known AMR markers/mutations (Table 1). The SM DNA extraction method was selected over the SP method for this study based on speed (1.5 hours time saving). Results from SP extracted gDNA were comparable in quality and quantity (data not shown). *B. anthracis* SM extractions produced HMW DNA (>30 kb) with minimal shearing (Supp. Fig. 7) at concentrations acceptable for MinION sequencing (Supp. Table 1). The SM extraction for *B. anthracis* Sterne/pUTE29 met ONT’s absorbancy ratio requirements, but the A260/230 ratio for strains 411A2 and UT308 were lower than the recommended range of 2.0-2.2. Repeated extraction attempts did not improve the ratios for these strains (data not shown). Additional clean-up by DNA precipitation was not performed to avoid extending the procedure time and to minimize gDNA shearing. Rapid Sequencing Kit libraries were prepared using gDNA from UT308, 411A2, and Sterne/pUTE29 and Field Sequencing Kit libraries were prepared using gDNA from the latter two strains.

### *B. anthracis* MinION WGS and retrospective down sampling analysis

Following nanopore sequencing of *B. anthracis* strains, 100,000 passed fast5 reads were analyzed (Fig. 5). The average number of mismatches among the Rapid and Field Sequencing Kit *B. anthracis* data sets ranged from 169 to 594 (Fig. 5a). *B. anthracis* Sterne/pUTE29 Illumina MiSeq and PacBio assemblies contained fewer mismatches (107 and 87, respectively). Nanopore *B. anthracis* data sets contained thousands of indels (6,262-11,574), compared to Illumina (24 indels) and PacBio (21 indels) assemblies (Fig. 5b). Nanopolish increased the fraction of *B. anthracis* poly-N lengths correctly called, but this fraction decreased with lengths >4. Increased accuracy was observed when calling poly-As and -Ts compared to poly-Gs and -Cs (Supp. Fig. 8).

**Figure 5.**
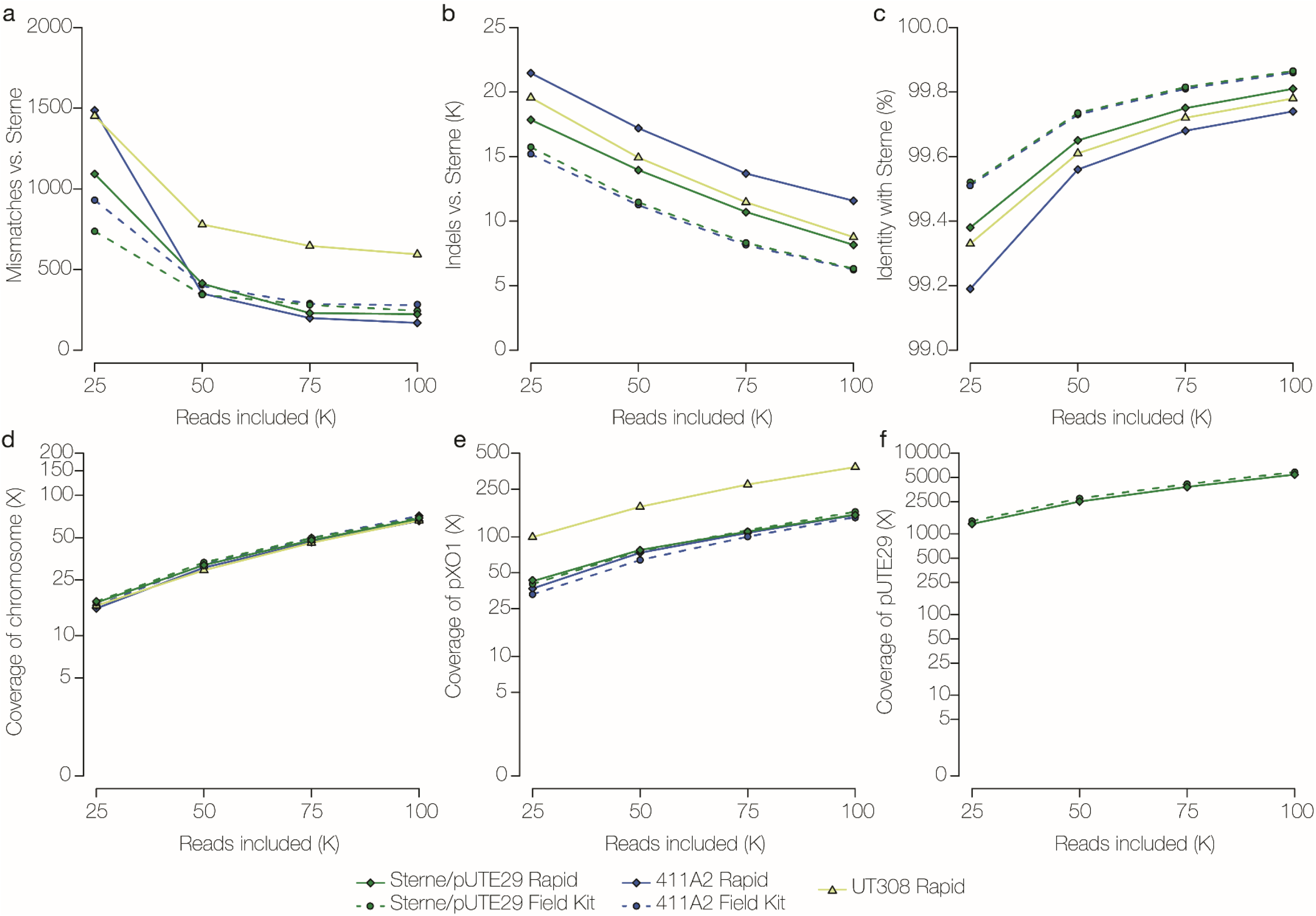
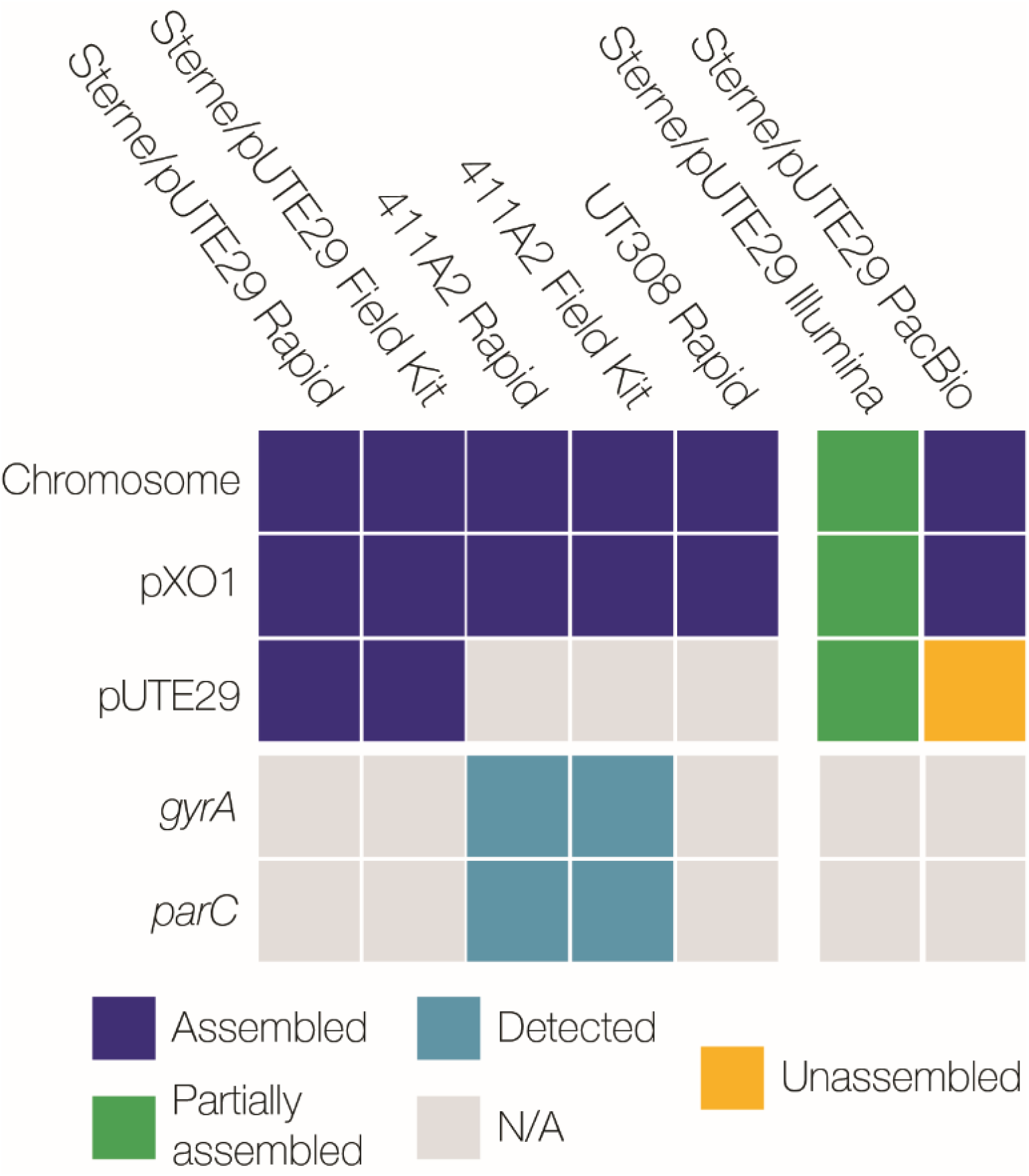
Assessment of nanopore sequencing performance and *de novo* assembly for *B. anthracis* strains. DNA sequencing libraries were prepared using the Rapid and Field Sequencing Kits from a silica-membrane extraction. Assemblies were generated using analysis of the first 25,000, 50,000, 75,000, and 100,000 passed fast5 nanopore reads. **a-c** Determination of mismatches, indels and % identity, and **d-f** fold coverage for chromosomal and plasmid sequences in data sets of *B. anthracis* strains (Sterne/pUTE29, 411A2, and UT308) aligned to the Ames Ancestor reference sequence. **g** Using analysis of 100,000 passed fast5 reads, the ability to assemble the chromosome and plasmids (blue) and detect the *gyrA* and *parC* SNPs associated with the non-susceptible CIP phenotype (teal) or wild type sequence (N/A), is depicted in a presence/absence plot. Assembled is defined as a complete sequence with the same nucleotide order as the reference sequence (navy), partially assembled represents a sequence that was assembled but not as a single contig (green), and unassembled refers to a sequence that was not present in the assembly (orange).

The 100,000-read analysis yielded >99% identity across the ~5.2 Mb *B. anthracis* genome in every data set analyzed (Fig. 5c). *B. anthracis* nanopore assemblies contained single contigs for the chromosome (66X average depth of coverage) and pXO1 (~180 kb, 223X average depth of coverage), which were consistantly assembled to completion (Fig. 5d, e, g). Analysis of 100,000 fast5 reads provided sufficient coverage to detect known AMR markers (chromosomal or plasmid-associated) (Table 1). For Sterne/pUTE29, the ~7.3 kb pUTE29 plasmid was consistantly assembled using both Rapid and Field Kit nanopore data (average 5,621X depth of coverage) (Fig. 5f, g). However, pUTE29 was partially assembled or unassembled using Illumina and PacBio data, respectively (Fig. 5g). The *gyrA* and *parC* mutations associated with quinolone resistance were detected in the CIP-NS strain 411A2 (Fig. 5g). Among the SM Rapid and Field Kit runs, the average time to generate 100,000 fast5 reads was ~3 hours (Supp. Table 2) and data analysis required ~ 2.5 additional hours (including base-calling). For *B. anthracis*, SM DNA extractions from an agar culture isolate, MinION WGS, and *de novo* assembly and analysis can be completed in ≤ 8.5 hours (Fig. 3).

## Discussion

During an outbreak or public health emergency involving bacterial biothreat pathogens, critical decisions are made immediately after a release, when details about the causative agent(s) are limited, and at every stage of the response. Rapid reporting of accurate laboratory testing results (pathogen identification, strain typing, AST) are critical for informing a public health response. For example, detection of genetic manipulation and/or unexpected AMR profiles could signal the introduction of plasmids, AMR genes, or other evidence of engineering. Speeding this response to limit the impact of biothreat incidents is a goal of the 2018 U.S. National Biodefense Strategy^39^

Culture- and DNA-based methods have improved the ability of laboratories to rapidly characterize microbial pathogens. The introduction of NGS platforms has accelerated these efforts, and WGS offers a comprehensive genome-wide snapshot of microbial pathogens. However, DNA sequencing using Illumina and PacBio platforms is time-consuming compared to nanopore sequencing and relies on instuments with a large footprint that are often space-prohibitive. Advances in long-read microbial DNA sequencing with the MinION instrument have transformed the fields of NGS and molecular characterzation, enabling real-time pathogen sequencing for epidemiologic investigations^40^. MinION WGS is promising as a front-line strategy for on-site microbial characterization due to the speed of sample preparation, relatively low cost and device portability^41^. Here, we describe a nanopore sequencing approach for *Y. pestis* and *B. anthracis* with a turnaround time that is ~90% faster than current Illumina MiSeq workflows and yields complete genome assemblies, including large and small plasmids, and accurately identifies AMR genes and mutations within a work day.

For the widespread adoption of WGS-based approaches in public health and clinical laboratories, validated methods that are simple to set up are essential. During an outbreak investigation, laboratory methods must be reliable and reproducible. Both commercially-available DNA extraction methods evaluated here were capable of generating gDNA of sufficient quality, quantity, and molecular weight for *Y. pestis* and *B. anthracis* MinION sequencing. The SM method was the fastest (30 min for *Y. pestis* and 1.5 hr for *B. anthracis*) and simple to perform. Despite the sub-optimal quality of some *B. anthracis* extractions, *de novo* assemblies using data from these samples were unaffected, indicating downstream library preparation and sequencing reactions were not compromised. Further optimization could improve the extraction process and DNA quality, and work is ongoing to evaluate the sequencing performance of DNA extracted from wild type *B. anthracis* encapsulated vegetative cells.

Shelf-stable reagents that do not require cold chain storage are preferred for rapid deployment and use in the field. The lyophilized Field Sequencing Kit, which does not require a cold chain, performed comparably to the Rapid Kit. To identify the minimum amount of sequencing time needed to consistently generate enough data for high-quality *Y. pestis* and *B. anthracis de novo* genome assemblies, a retrospective down sampling analysis was performed. On average, only 100,000 fast5 reads were necessary to assemble the chromosome and large and small plasmids with sufficient coverage to correctly identify known AMR genes and mutations. Additional reads did not strengthen the assembly quality, and any improvement of coverage and number of mismatches, indels, and % identity was negligible. Our laboratory and bioinformatic workflows were achieved in <8.5 hours starting from a culture isolate, and accurately detected genomic AMR markers in approximately half the time required to complete functional AST for *Y. pestis* and *B. anthracis*.

We demonstrate that thresholds and quality metrics can be established that are indicative of nanopore sequencing quality. MinION sequencing performance was assessed by comparing data from two technical replicate experiments where the DNA extraction, library preparation kit, sequencing conditions, MinKNOW software, and MinION instrument and flowcell versions remained the same. Since the majority of nanopore data is generated during the first 16 hours of sequencing when the most active pores are used^26, 36^, only the passed reads generated during this time were analyzed in order to standardize the technical replicate data sets. Considerable run-to-run variation was observed among the SP and SM replicates from the same biological sample. We demonstrated that the number of active pores was not predictive of the amount or quality of data. These inconsistencies could result from intrinsic flow cell variability and day-to-day loading efficiencies. However, comparable average quality scores were observed for basecalled reads (≥Q7) for both the SP and SM data sets.

The rapid sequencing workflow developed in this study aimed to deliver the fastest turnaround time per bacterial isolate using one flowcell per run. The study avoids the complexities of multiplexing/demultiplexing sequences during data analysis. Due to the variability we observed in flowcell performance, multiplexing the same sample on a single flow cell could be used to establish “within run” precision. Improved rapid multiplex capabilities have been described since this study began, but multiplexing is a balance between sequencing speed and genome coverage. Additional considerations, including incorporation of positive and negative DNA controls, are needed to establish indicators of sequencing quality within and between runs. Future work must build on this foundation to ensure the accuracy of nanopore data and expand MinION sequencing to include multiple complex samples, such as clinical specimens.

The assay parameters established in the *Y. pestis* technical replicate analysis were applied to *B. anthracis* by sequencing three AMR strains: Sterne/pUTE29, 411A2, and UT308. With the exception of an additional one hour lysis step for *B. anthracis*, the same SM DNA extraction protocol was employed and 100,000 fast5 reads were sufficient for >99% coverage of the chromosome and plasmids (both native and introduced). While functional phenotypic methods remain essential for accurate AST, WGS can be used to detect point mutations that predict AMR in some biothreat pathogens^42^ For both *Y. pestis* and *B. anthracis* data sets, MinION reads were error-prone compared to Illumina and PacBio data and contained more indels, a well described nanopore error type^33, 34, 43^. Therefore, open reading frame prediction and SNP-typing were not performed for MinION assemblies due to the large number of indels. Overall, use of the Nanopolish error correction tool reduced indels, but accurate basecalls could not be made in homopolymer regions (poly-N lengths >4). For this reason, the homopolymeric regions associated with penicillin resistance in *B. anthracis*, *sigP-rsiP*^42^, were not included for analysis in this study. However, QRDR mutations associated with AMR in *B. anthracis* and *Y. pestis* that are located in non-homoploymeric regions of the genome were correctly identified in nanopore assemblies. Currently, a hybrid (long- and short-read) assembly approach is necessary to construct complete genomes with sufficent quality for variant calling (for SNP and indel detection) with nanopore data.

*Y. pestis* isolates harboring various multimeric pPCP1 forms in addition to, or in place of, one of the three prototypical plasmids were described previously^38, 44^ The 9.5 kb and 19 kb pPCP1 plasmids are stable upon multiple passages and coexist; however, one plasmid is often present at a higher concentration than the other, suggesting a level of genetic control of expression^38^. Chu *et al*.^38^ noted the presence of pPCP1 dimer forms in strain A1122, regardless of DNA extraction method used^38^. Other atypical *Y. pestis* plasmids have been described in Angola and C790 strains, including a chimeric 114,570 nt form with two tandemly repeated pPCP1 copies integrated into pMT1^45^, and a pMT1-pCD1 chimera^46^, respectively. The presence of an IS element, IS*100*, on pPCP1, pCD1, and pMT1 may have facilitated the formation of these multimeric and chimeric plasmids via homologous recombination^45, 46^. Understanding these plasmid size and form variations among naturally occurring *Y. pestis* isolates is important for distinguishing between natural and potentially engineered variation^44^

In this study, the pPCP1 monomer and dimer forms were not consistently assembled correctly because multiple pPCP1 reads of different sizes introduced considerable ambiguity into the assembly graphs. Multiple overlapping pPCP1 reads corresponding to closed-circle multimer plasmids of various increasing sizes were also identified. The longest pPCP1 multimers were detected in a *Y. pestis* S1037 assembly for which additional steps to reduce gDNA shearing during the SP extraction were employed. Analysis of 100,000 reads revealed four 5-mer reads and one 6-mer read. Additional analysis of 300,0 reads identified eleven 5-mer reads and only one 6-mer read (data not shown); none of which were detected in SP technical replicate data sets. The challenges of using long-read sequence data for plasmid assembly, discussed below, and/or DNA shearing may explain the rarity of finding these large multimers.

Chu *et al*.^38^ also described the presence of multimeric pPCP1 forms in A1122 >19 kb by Southern blot analysis and they explain these multimer forms may contribute to bacterial plasmid maintenance. While the *Y. pestis* genome is considered monomorphic^47^, there is evidence for genome plasticity due to large genomic rearrangements (translocations and inversions), as well as gene loss due to homologous recombination between multiple copies of highly repetitive IS elements found on all three plasmids and on the chromosome^48, 49, 50^. We speculate that in *Y. pestis* strains S1037 and All22 the 9.5 kb pPCP1 plasmid has undergone homologous recombination with itself and with the 19 kb pPCP1 plasmid to form the 2-mer to 6-mer species.

The power of MinION sequencing to resolve complex genomic structures was evident in the ability to identify expected and novel pPCP1 plasmid multimers. These pPCP1 multimers were only partially assembled using S1037 and A1122 Illumina and PacBio data. In addition, the A1122 reference sequence (CP002956.1), assembled from Illumina data, contains only the pPCP1 monomer form. This is an example of the limitation of using short-read data to assemble complex genomic regions and highlights that assemblies in public databases may be incomplete.

Small plasmid assembly using long-read data is challenging for many algorithms^51^. Unlike the issues associated with assembling multiple copies of the *Y. pestis* multimeric plasmid, the pUTE29 (~7.3 kb) plasmid was readily assembled in all *B. anthracis* data sets. However, the bioinformatics pipeline in this study was tailored to assemble these plasmids. It is important to note that these parameters may not be generalizable for every data set. While this work was performed using a small number of strains, the availablity of resistant *Y. pestis* and *B. anthraicis* isolates is limited; for both laboratory derived and naturally-occuring isolates. Further optimization is required to assure the pipeline can correctly identify different plasmid and AMR profiles, and this is currently under investigation. Since the completion of this work, new assemblers, e.g. WTDBG2^52^ and FLYE^53^, have been released and address the limitations of plasmid assembly using long-read sequence data.

Other limitations of nanopore sequencing (e.g. cost, throughput, and accuracy) could be mitigated by future updates to the software, hardware, library preparation kits, and reagents^41^. But, with these improvements and updates come challenges such as locking down assay parameters to assess and compare performance^36, 54, 55^. The workflows described here for same-day bacterial MinION WGS, assembly, and bioinformatics analysis for biological threat pathogens show the utility of nanopore sequencing for rapid diagnostics that will be critical during a biothreat event. As NGS technologies continue to evolve and stablize, real-time nanopore WGS for bacterial characterization, including AMR marker detection, chromosome, large and small plasmid assembly, and accurate SNP detection, may soon be a reality.

## Methods

### Biosafety Procedures

Only select agent-excluded strains were used in this study (Y. *pestis* A1122, *Y. pestis* S1037, *B. anthracis* Sterne/pUTE29, *B. anthracis* 411A2, and *B. anthracis* UT308). All procedures were performed in a biosafety level 2 (BSL-2) laboratory by trained personnel wearing appropriate personal protective equipment (PPE) according to the CDC/NIH publication *Biosafety in Microbiological and Biomedical Laboratories*, 5th edition^56^.

### Bacterial Strains

Study strains are listed in Table 1. *Y. pestis* A1122 was passaged on increasing concentrations of CIP using a previously described method^11^ to create the CIP-NS strain, S1037. This strain was developed as a control strain with approval from the CDC Institutional Biosafety Committee and Institutional Biosecurity Board.

### Growth Conditions

All strains were cultured for 16-20 hours (*B. anthracis*) or 24 hours (*Y. pestis*) on BD BBL^TM^ Trypticase Soy Agar II with 5% sheep blood (SBA) (Thermo Fisher Scientific, Waltham, MA, USA) at 35°C in ambient air from glycerol stocks stored at −170°C.

### Antimicrobial Susceptibility Testing

Broth microdilution (BMD) was performed to determine antimicrobial susceptibility following the Clinical and Laboratory Standards Institute guidelines^57^ Genomic DNA Extraction. gDNA for each study strain was extracted from agar plate cultures, described above. For *Y. pestis*, colonies were transferred to 200μl of Phosphate Buffer Solution (0.01M, pH 7.40) and vortexed until completely resuspended. *B. anthracis* colonies were transferred to 200μl of lysis buffer (20mM Tris HCL pH 8.0, 2mM EDTA, 1.2% Triton X-100) and incubated for 60 min at 37°C. *Y. pestis* and *B. anthracis* gDNA was isolated using the MasterPure Complete DNA and RNA Purification Kit (Lucigen, Middleton, WI), a salt-precipitation (SP) extraction method, and the QIAamp DNA Blood Mini Kit (Qiagen Valencia, CA), a silica-membrane (SM) extraction method. Both extractions were performed according to manufacturer’s specifications. *Y. pestis* extracts were eluted in 75μl of Elution Buffer (EB) (10 mM Tris-Cl, pH 8.5, Qiagen, Valencia, CA) and *B. anthracis* extracts were eluted in 50μl EB. gDNA samples were subsequently stored at 4°C.

### Assessment of gDNA quality and quantity

gDNA extractions were evaluated for quantity and quality using the Qubit dsDNA BR Assay Kit (Invitrogen, Waltham, MA) and NanoDrop 2000 spectrophotometer (ThermoFisher Scientific, Pittsburgh, PA) following manufacturer’s instructions. gDNA integrity was assessed following 0.6% agarose gel electrophoresis of 100ng of each sample to visualize the presence of intact, high molecular weight (HMW) DNA. The Quick-Load 1 kb Extend DNA Ladder (New England BioLabs, Ipswich, MA) was used as a molecular marker.

### Nanopore DNA library preparation

Libraries were prepared using 400ng of gDNA per strain and the Rapid Sequencing Kit, SQK-RAD004 (ONT, Oxford, England) or the Field Sequencing Kit, SQK-LRK001 (ONT, pre-released), according to manufacturer’s specifications. All Field Sequencing Kit components were stored at room temperature prior to use. *Y. pestis* S1037 libraries were prepared in triplicate on the same day from the same gDNA extraction and stored at 4°C for no longer than 3 days.

### Nanopore sequencing

Sequencing was performed on a MinION (MK1B) for 16-48 hours using R9.4.1 flowcells and MinKNOW (version 18.01.6). All flowcells met the minimum ≥800 active pore count requirement immediately prior to sequencing. Each sample was sequenced on a separate flowcell and each flowcell was used for only one sequencing run. For the technical replicate study, the same laboratorian loaded each library into the flowcells to minimize variation. The replicate *Y. pestis* S1037 libraries were sequenced on sequential days over a 3 day period. To standardize the comparison of the technical replicate Rapid Kit and Field Kit *Y. pestis* S1037 data sets, only raw fast5 reads from the “passed folder,” generated during the first 16 hours of sequencing were analyzed. Raw fast5 reads from the “failed folder” were not included in this analysis. Sequencing runs included in this study are described in Supp Table 2.

### Bioinformatics pipeline

A pipeline was developed to perform rapid *de novo* genome and plasmid assemblies, to detect AMR variants and genes, and to identify variants (mismatches and indels), see Supp Fig. 9.

### Basecalling

Basecalling of raw fast5 reads was performed using the Albacore basecalling utility (version 2.3.10), to produce raw FASTQ reads. Raw FASTQ reads were further processed using Porechop (version 0.2.3) to remove adapters and any potentially chimeric sequences. Processed FASTQ reads were used to generate a draft assembly using minimap2^58^ (version 2.10-r770-dirty) and miniasm^59^ (0.2-r168-dirty). First, all-versus-all alignment of the FASTQ reads was performed using minimap2 with the option ‘-x ava-ont’. The resulting minimap2 alignments and input reads were used to create a draft assembly with miniasm, using the options ‘-s 1750 –h 1000 –1.5’. Errors in the draft assembly generated by miniasm were iteratively improved using RACON^60^ (version 1.1.1). For 5 iterations, minimap2 was used to align the reads to an assembly and the alignments were used by RACON to produce a consensus assembly. The first iteration used the draft assembly generated by miniasm, while subsequent iterations used the consensus assembly generated by the previous RACON step. The final output of the RACON process was further processed using Nanopolish^61^ (version 0.10.1). Nanopolish was run in the variants mode using the options ‘--faster --consensus --min-candidate-frequency 0.1’, and the vcf2fasta feature was used to create the final consensus sequence.

### Detection of known genes related to AMR

The ResFinder^62^ database of AMR genes was queried against the final Nanopolish assembly using BLASTn^63^ and the options ‘-perc_identity 90 – evalue 1e-10’ to identify potential AMR sequences in each assembly. AMR gene results were clustered using BEDTools^64^ merge (version 2.24.0) with the options ‘-d −30’ in order to identify unique, nonoverlapping regions with high similarity to AMR sequences. For each cluster, BEDTools intersect was used to find the gene with the highest % identity, which most completely covered the cluster.

### Detection of known mutations related to AMR

Basecalls from the nanopore sequencing reads were compared to a reference genome using minimap2^58^ by aligning reads against the corresponding reference sequence (*Y. pestis* A1122, Accession No. CP002956.1, or *B. anthracis* Ames Ancestor, Accession No. AE017334) with the options ‘-a -x map-ont’. The samtools^65^ mpileup utility (version 1.2) was used to find the numbers of each base seen in the reads at each position in the QRDR gene coding regions. These basecall counts were used with bcftools^65^ call (version 1.8) to identify variant sites in those genes.

### Characterization of Plasmids

Contigs ≥500,000 nt were excluded from plasmid characterization as they likely represent chromosomal sequence. Remaining contigs (≤500,000 nt) were queried with a custom database of plasmids and artificial sequences using the pblat^66^ utility (version 35). A custom algorithm, pChunks, was used to find the set of plasmid sequences present in the assembly such that the maximal amount of plasmid sequence was included.

### Detection of mismatches, insertions and deletions

The consensus assembly generated by the last iteration of RACON was aligned with the corresponding reference sequence (Y. *pestis* A1122 or *B. anthracis* Ames Ancestor) to identify mismatches and small indels using the dnadiff utility from the MUMmer^67^ package (version 4.0.0beta2). Gaps from the resulting ‘rdiff (reference differences) and ‘qdiff (query differences) outputs were used to identify indels in one genome relative to the other. Sequence gaps ≥100 bases, and with positive lengths in both the assembly and reference sequences (denoting real insertions and not duplications) were kept as insertions. Mismatches identified by MUMmer were compared to the presence of homopolymeric regions in the reference sequence to determine the number of homopolymeric regions that were assembled correctly. Coverage of the reference chromosome and plasmids was characterized by aligning reads against the references sequences using minimap2, and enumerating the mean per-base coverage.

### Downsampling

*Y. pestis* and *B. anthracis* reads were retrospectively assembled using the above analysis in the order they were generated for the first 25,000, 50,000, 75,000, 100,000 passed fast5 reads to determine the minimum number of sequencing reads required for chromosome and plasmid assembly, as well as the identification of known AMR genes and mutations. Per assembly, the number of mismatches, indels, and percent identity were determined using an alignment to the reference sequence. The chromosome and plasmid precent coverage was calculated.

### pPCP1 Analysis

For each SP S1037 *Y. pestis* 100,000-read data set, basecalled reads of ≥1000 nt were aligned to the pPCP1 reference sequence from *Y. pestis* A1122, Accession No. CP002956.1, using minimap2. Reads that aligned to the pPCP1 sequence across at least 95% of their length were included for analysis. Reads were assigned to a 1, 2, 3, 4, 5, or 6-mer version of the pPCP1 sequence based on their length and the fractions of these versions in each sample were used to determine the pPCP1 multimer fraction (Supp Fig. 4). All SP *Y. pestis* S1037 data sets contained reads corresponding to 2, 3, and 4-mer forms of plasmid pPCP1, and SP Rapid 1 was chosen as the representative data set to use to generate Circos plots^68^ representing the distribution of reads from the plasmid multimers. From the SP Rapid 1 aligned reads, a representative read for each multimeric pPCP1 sequence was choosen to represent the complete plasmid. The representative read was rotated such that the first base of the read aligned to the first base of the pPCP1 reference sequence. The remaining reads for each multimeric pPCP1 sequence were aligned against the reference read. Known coding sequences from pPCP1 were found in the multimeric forms by querying these features against the representative read using BLASTn^63^.

### Illumina library preparation, sequencing, and analysis

Sequencing of *Y. pestis* S1037 and *B. anthracis* Sterne/pUTE29 was performed on an Illumina MiSeq using the v2 kit (2×250 nt paired-end) with Nextera XT libraries. Sequencing adapters were removed and reads were trimmed using Cutadapt (v1.14). *De novo* assembly was performed using the SPAdes Genome Assembler (v3.10.0). The consensus Illumina assemblies were aligned with the corresponding *Y. pestis* A1122 or *B. anthracis* Ames Ancestor reference assemblies to identify mismatches and small indels using the dnadiff utility from the MUMmer^67^ package (version 4.0.0beta2).

### PacBio library preparation, sequencing, and analysis

Libraries (10 kb) of *Y. pestis* S1037 and *B. anthracis* Sterne/pUTE29 were prepared with the SMRTbell Template Prep Kit 1.0 (Pacific Biosciences, Menlo Park, CA) and bound using the DNA/Polymerase Binding Kit P6v2 (Pacific Biosciences, Menlo Park, CA). The bound libraries were then loaded on one SMRTcell and sequenced with C4v2 chemistry for 270 min movies on the RSII instrument (Pacific Biosciences, Menlo Park, CA). Sequence reads were *de novo* assembled using RS_HGAP Assembly.3 in the SMRT Analysis 2.3.0 portal. The consensus PacBio assemblies were aligned with the corresponding *Y. pestis* A1122 or *B. anthracis* Ames Ancestor reference assemblies to identify mismatches and small indels using the dnadiff utility from the MUMmer^67^ package (version 4.0.0beta2). Comparison of PacBio reads corresponding to the pPCP1 sequence was carried out as described above for the analysis of pPCP1.

## Data Availability

The nanopore, illumina, and PacBio sequence data sets generated and analyzed in the current study are available from NCBI in the Sequence Read Archive under bioproject PRJNA523610. All nanopore sequencing runs and corresponding BioSample accession numbers are also listed in Supp Table 2. All data supporting the findings of this study are available within the article and its supplementary information files, or are available upon request.

## Acknowledgements

This work was supported by the U.S. CDC Office of Public Health Preparedness and Response (now Center for Preparedness and Response). We thank Christopher Lynberg, CDC, and Charles Johnston, General Dynamics contract agency, for computational support, technical advice, and valuable discussions. We thank Dhwani Batra, CDC, and Lori A. Rowe, CDC, for PacBio sequencing and analysis. We thank Jeannine Petersen, CDC, Luke C. Kingry, CDC, and Alex Hoffmaster, CDC, for their critical review of the manuscript. The findings and conclusions in this report are those of the authors and do not necessarily represent the views of the Centers for Disease Control and Prevention. The use of trade names is for identification only and does not imply endorsement by the Centers for Disease Control and Prevention.

## Author Contributions

A. S.G. and D.S. conceived the project and all authors contributed to the design of experiments. A.S.G., B. C., and H.P.M. performed all experiments and all authors contributed to data analysis. A.B.C. established the bioinformatics pipeline, performed all sequence analysis, and created all figures. A.S.G. led development of the manuscript, and all authors contributed to the writing process. All authors reviewed and edited the manuscript.

