## Supplemental Figures and Tables for "Rapid Detection of Genetic Engineering, Structural Variation, and Antimicrobial Resistance Markers in Bacterial Biothreat Pathogens by Nanopore Sequencing"

### Supplementary Material

**Supplemental Table 1**. *Y. pestis* and *B. anthracis* DNA extraction quality and quantity.

| **Strain** | **Extraction** | **A 260/280nm** | **A 260/230nm** | **Estimated DNA Concentration (ng/µl)** |
|---|---|---|---|---|
| *Y. pestis* A1122 | SP | 1.94 | 1.78 | 69.7 |
| *Y. pestis* S1037 | SM | 1.97 | 2.11 | 81.5 |
| | SP-1 | 1.95 | 2.06 | 67 |
| | SP-2 | 1.91 | 2.15 | 78 |
| *B. anthracis* Sterne/pUTE29 | SM | 1.96 | 2.16 | 103 |
| *B. anthracis* 411A2 | SM | 1.87 | 1.53 | 60.9 |
| *B. anthracis* UT308 | SM | 1.96 | 1.56 | 81.8 |

(A) absorbance, (SM) silica-membrane extraction, (SP) salt-precipitation extraction

**Supplemental Table 2**. Details of *Y. pestis* and *B. anthracis* nanopore sequencing runs included in this study.

| ***Y. pestis*** **Strain (Extraction)** | **Library Preparation** | **Run Date** | **MinKNOW Sequencing Script** | **Total Run Time** | **Time to 100,000 passed fast5 reads** | **Strand to Pore Ratio (~10 run time)** | **fast5 Accession Number** | **FASTQ Accession Number** |
|---|---|---|---|---|---|---|---|---|
| S1037 (SM) | Rapid 1 | 2/14/2018 | SQK-RAD004 | 17 hr 54 min | 1 hr 2 min | 263/326 (80%) | SRR8608868 | SRR8608876 |
| S1037 (SM) | Rapid 2 | 2/15/2018 | SQK-RAD004 | 17 hr 1min | 57 min | 239/317 (75%) | SRR8608869 | SRR8608877 |
| S1037 (SM) | Rapid 3 | 2/16/2018 | SQK-RAD004 | 24 hr 9 min | 1 hr 38 min | 256/312 (82%) | SRR8608870 | SRR8608878 |
| S1037 (SM) | Field | 03/14/2018 | SQK-RAD003 | 28 hr 27 min | 1hr 35 min | 298/340 (87%) | SRR8608871 | SRR8608879 |
| S1037 (SP-1) | Rapid 1 | 2/28/2018 | SQK-RAD004 | 20 hr 29 min | 1 hr 37 min | 153/252 (60%) | SRR8608853 | SRR8608872 |
| S1037 (SP-1) | Rapid 2 | 3/1/2018 | SQK-RAD004 | 23 hr 29 min | 2 hr 41 min | 159/251 (63%) | SRR8608852 | SRR8608873 |
| S1037 (SP-1) | Rapid 3 | 3/2/2018 | SQK-RAD004 | 46 hr 24 min | 3 hr 18 min | 157/244 (64%) | SRR8608855 | SRR8608874 |
| S1037 (SP-1) | Field | 3/5/2018 | SQK-RAD003 | 24 hr | 1 hr 57min | 257/344 (74%) | SRR8608854 | SRR8608875 |
| S1037 (SP-2) | Rapid | 09/18/2018 | SQK-RAD004 | 43 hr 47min | 8 hr 12 min | 84/218 (38%) | SRR8608849 | SRR8608880 |
| A1122 (SP) | Rapid | 09/19/2018 | SQK-RAD004 | 49 hr 16 min | 1 hr 47min | 260/347 (74%) | SRR8608851 | SRR8608862 |
| ***B. anthracis*** **Strain (Extraction)** | **Library Preparation** | **Run Date** | **MinKNOW Sequencing Script** | **Total Run Time** | **Time to 100,000 passed fast5 reads** | **Strand to Pore Ratio (~10 run time)** | **fast5 Accession Number** | **FASTQ Accession Number** |
| Sterne/pUTE29 (SM) | Rapid | 4/12/2018 | SQK-RAD004 | 24 hr | 3 hr 55 min | 107/201 (53%) | SRR8608850 | SRR8608863 |
| Sterne/pUTE29 (SM) | Field | 09/13/2018 | SQK-RAD003 | 40 hr 3 min | 1 hr 43 min | 208/330 (63%) | SRR8608847 | SRR8608864 |
| 411A2 (SM) | Rapid | 8/3/2018 | SQK-RAD004 | 48 hr | 1 hr 36 min | Not recorded | SRR8608846 | SRR8608865 |
| 411A2 (SM) | Field | 6/28/2018 | SQK-RAD003 | Not recorded | 4 hr 53 min | Not recorded | SRR8608860 | SRR8608866 |
| UT308 (SM) | Rapid | 6/27/2018 | SQK-RAD004 | 22 hr 47 min | 2 hr 15min | 87/237 (37%) | SRR8608861 | SRR8608867 |

All runs were sequenced using the Rapid Sequencing Kit (SQK-RAD004), or the Field Sequencing Kit (SQK-LRK001), R9.4.1 flow cells, and MinKNOW (Version 18.01.6). (SM) silica-membrane extraction, (SP) salt-precipitation extraction

**Supplemental Table 3**. Metrics of 16 hour nanopore sequencing for *Y. pestis* strain S1037.

| ***Y. pestis* S1037 Sequencing Run** | **Active Pores** | **Passed Reads** | **Failed Reads** | **≥Q7 Reads (Albacore)** | **# Reads Lost After Basecalling** | **% Reads Lost After Basecalling** | **Avg. Read Length (nt)** | **Longest Read (nt)** | **Data Generated (Gb)** | **Avg. Quality Score** |
|---|---|---|---|---|---|---|---|---|---|---|
| SP Rapid (1) | 1,061 | 676,497 | 54,793 | 639,598 | 36,899 | 5.45 | 4,769 | 144,371 | 3.1 | 13.1 |
| SP Rapid (2) | 888 | 410,515 | 46,382 | 388,435 | 22,080 | 5.38 | 4,690 | 144,371 | 1.8 | 13.2 |
| SP Rapid (3) | 759 | 329,875 | 85,554 | 310,906 | 18,969 | 5.75 | 8,239 | 126,776 | 2.6 | 13.1 |
| SM Rapid (1) | 1,358 | 1,190,946 | 75,040 | 305,824 | 885,122 | 74.32 | 4,377 | 101,096 | 1.3 | 12.6 |
| SM Rapid (2) | 1,276 | 1,294,029 | 144,023 | 1,172,803 | 121,226 | 9.37 | 3,927 | 149,903 | 4.6 | 12.5 |
| SM Rapid (3) | 968 | 650,090 | 180,273 | 477,864 | 172,226 | 26.49 | 5,533 | 93,179 | 2.6 | 12.8 |
| SM Field (1) | 1,326 | 621,837 | 52,943 | 594,145 | 27,692 | 4.45 | 5,030 | 86,368 | 3.0 | 13.9 |
| SP Field (1) | 1,502 | 579,770 | 60,298 | 545,442 | 34,328 | 5.92 | 6,284 | 116,662 | 3.4 | 13.5 |

**Supplementary Figure 1. Agarose gel electrophoresis of *Y. pestis* S1037 DNA.** gDNA (100 ng/lane) extracted using the silica-membrane (lane 2) and salt-precipitation (lane 3) methods was visualized. DNA Ladders (NEB 1 kb Extend, lanes 1 and 5; and NEB Supercoiled DNA Ladder, lane 6) and lambda phage gDNA (ONT, lane 4) were also included.

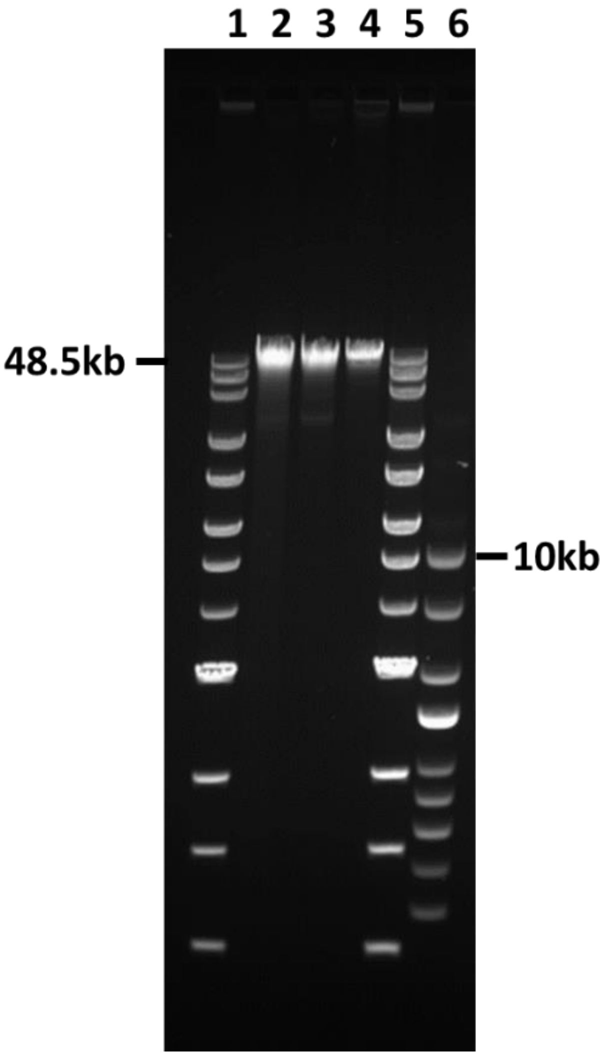

**Supplementary Figure 2. Poly-N length and fraction of poly-N's correct in *Y. pestis* S1037 assemblies using 100,000 fast5 read data sets. a** The average fraction of poly-A/T's or **b** poly-G/C's, with sequence lengths of 3 to 7 nucleotides, called correctly in *Y. pestis* S1037 nanopore assemblies using sequence polishing tools RACON (dotted line) and Nanopolish (solid line). The fraction correct was calculated as an average using all salt-precipitation and silica-membrane replicate data sets (n=6).

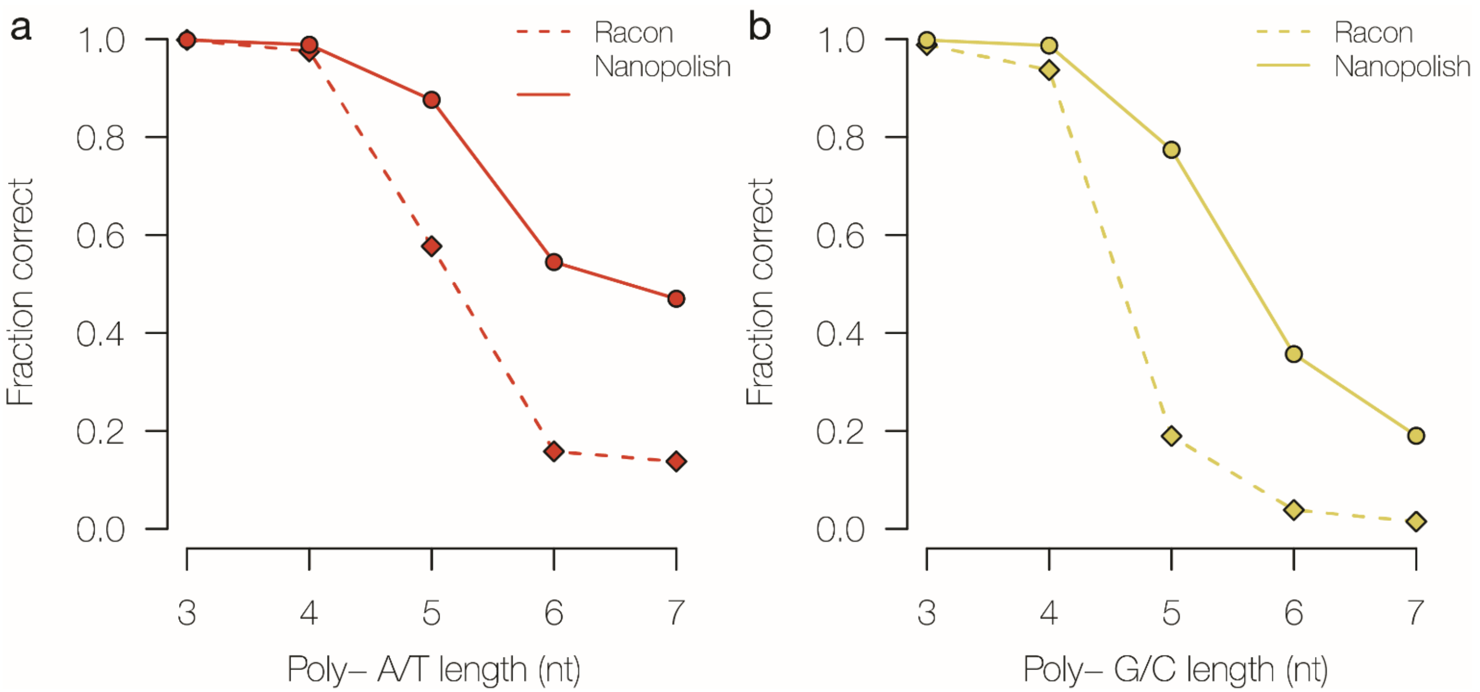

**Supplementary Figure 3. Distribution of nanopore sequencing reads by length using the first 100,000 fast5 nanopore reads analyzed for *Y. pestis* S1037 data sets. a-c** Overall read length distributions for the three salt-precipitation (SP) replicates. **d-f** Overall read length distributions for the three silica-membrane (SM) replicates. The peak in graphs a-c corresponds to the increased fraction of ~18.2 kb reads in the SP data sets that represent the pPCP1 dimer.

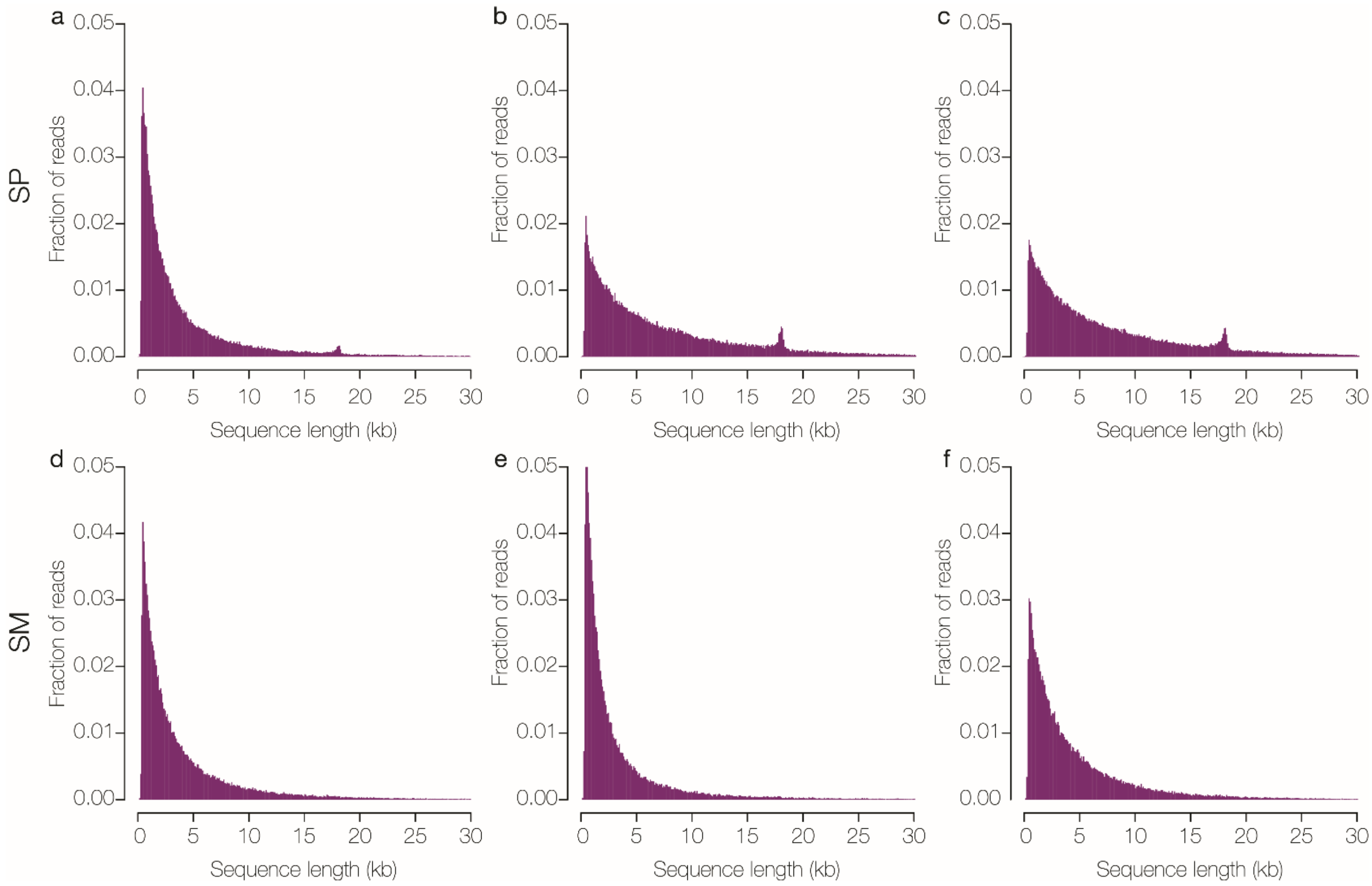

**Supplementary Figure 4. Fraction of the pPCP1 multimers found in *Y. pestis* S1037 nanopore data sets of 100,000 fast5 reads.** Based on length, reads were assigned to a multimeric form of pPCP1 (x-axis) and the fraction of each multimer is displayed (y-axis). Salt-precipitation (SP) data sets are in green and silica-membrane (SM) data sets are in purple. Each dot represents the multimer fraction of a rapid technical replicate and bars depict the average multimer fraction for all three technical replicates. The 3-mer and 4-mer forms were not detected in the SM data sets.

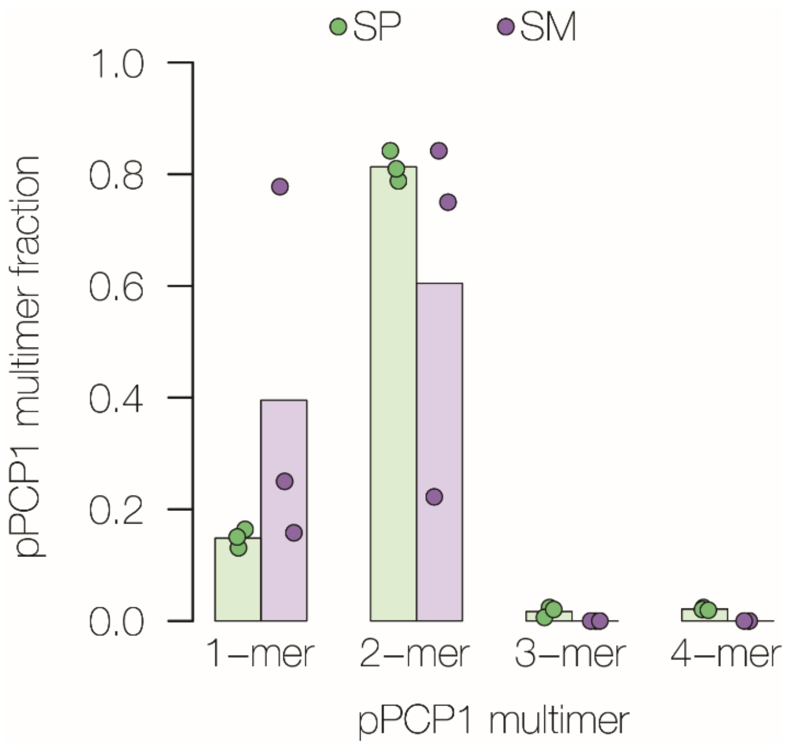

**Supplementary Figure 5. Distribution and fraction of reads in *Y. pestis* S1037 SP-2 100,000 read assembly. a** Distribution of nanopore sequencing reads by length using the first 100,000 fast5 reads analyzed for the *Y. pestis* S1037 SP-2 data set in which additional care was taken to minimalize shearing during extraction. The increased fraction of ~18.2 kb reads in the SP data sets corresponds to the pPCP1 dimer. **b** Fraction of nanopore sequencing reads with homology to pPCP1 using the first 100,000 fast5 reads analyzed for the *Y. pestis* S1037 SP-2.

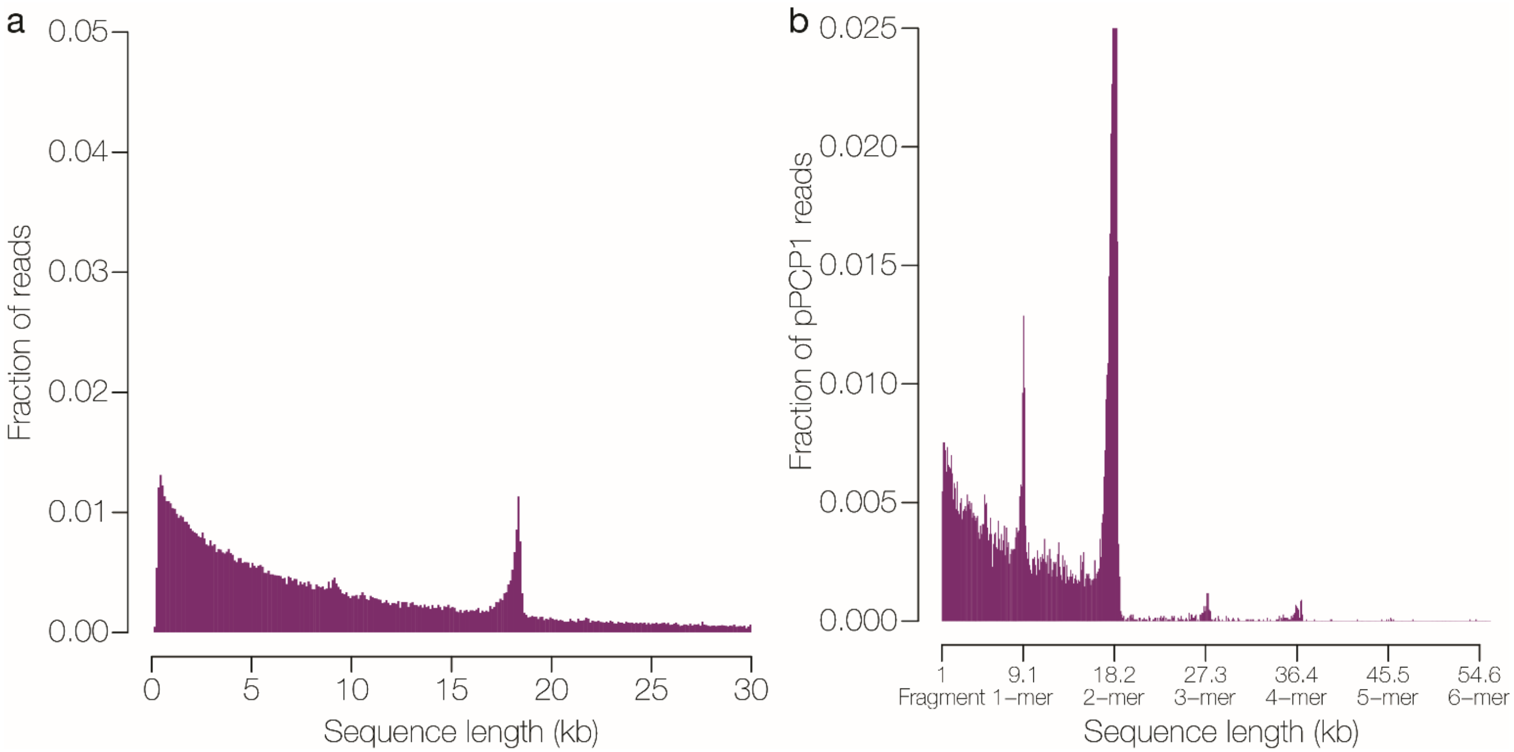

**Supplementary Figure 6. Distribution of nanopore sequencing reads by length using the first 100,000 fast5 reads analyzed for *Y. pestis* A1122**. **a** salt-precipitation (SP) data set, and **b** silica-membrane (SM) data set

.

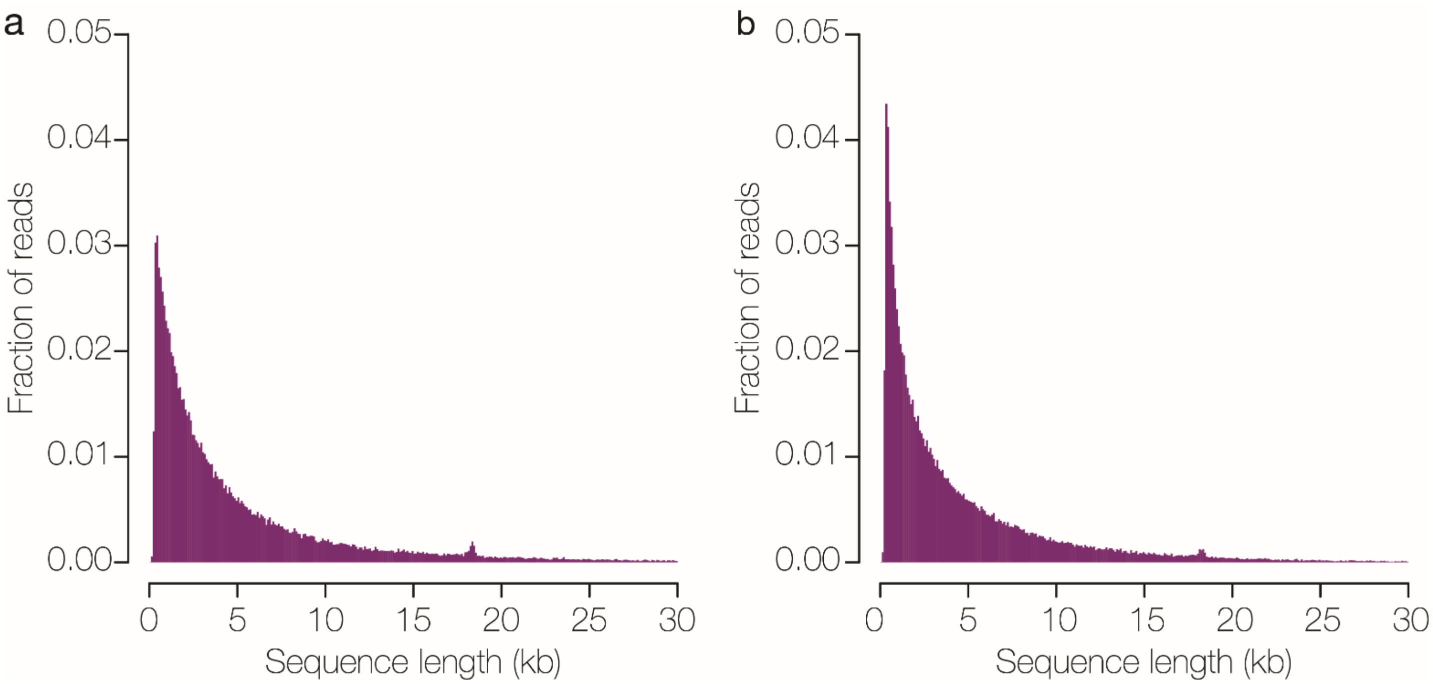

**Supplementary Figure 7. Agarose gel electrophoresis of *B. anthracis* DNA extracted using the silica-membrane method**. gDNA (100 ng/lane) was visualized for *B. anthracis* Sterne/pUTE29 (lane 2), 411A2 (lane 3), and UT308 (lane 4). DNA Ladder (NEB 1 kb Extend, lanes 1 and 6) and lambda phage gDNA (ONT, lane 5) were also included. The white line between lanes 2 and 3 indicates where the gel was cropped to remove lanes not relevant to this study.

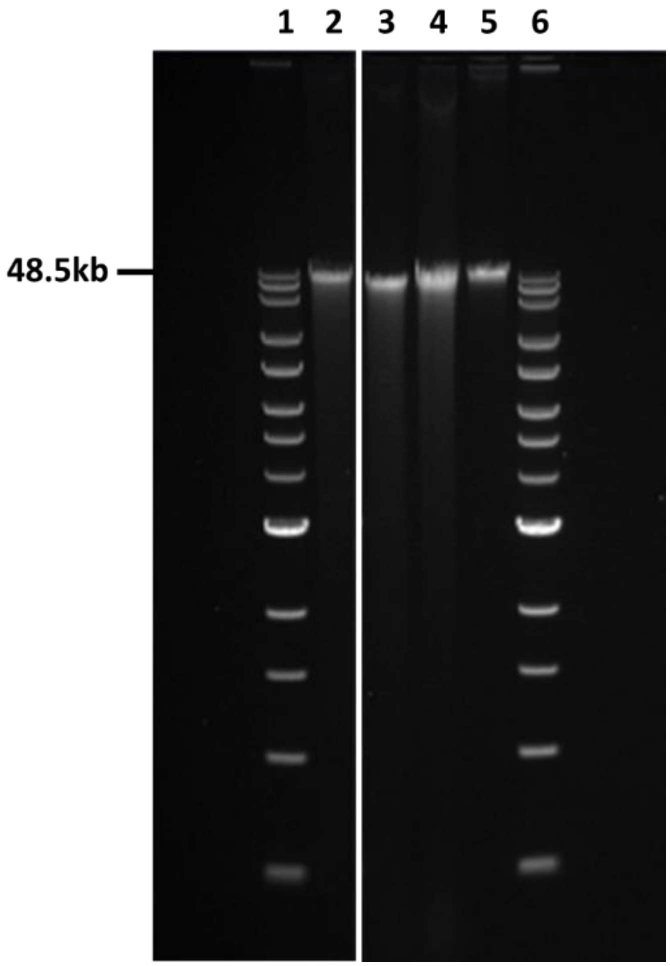

**Supplementary Figure 8**. **Poly-N length and fraction of poly-N's correct in *B. anthracis* Sterne/pUTE29 100,000 read assemblies. a** The average fraction of poly A/Ts or **b** poly G/C's, with sequence lengths of 3-7 nucleotides, called correctly in *B. anthracis* nanopore assemblies using sequence polishing tools RACON (dotted line) and Nanopolish (solid line). The 100,000 fast5 read data sets were used for this analysis and the fraction correct was calculated as an average using the *B. anthracis* Sterne/pUTE29 (Rapid and Field), 411A2 (Rapid and Field), and UT308 (Rapid) assemblies (n=5).

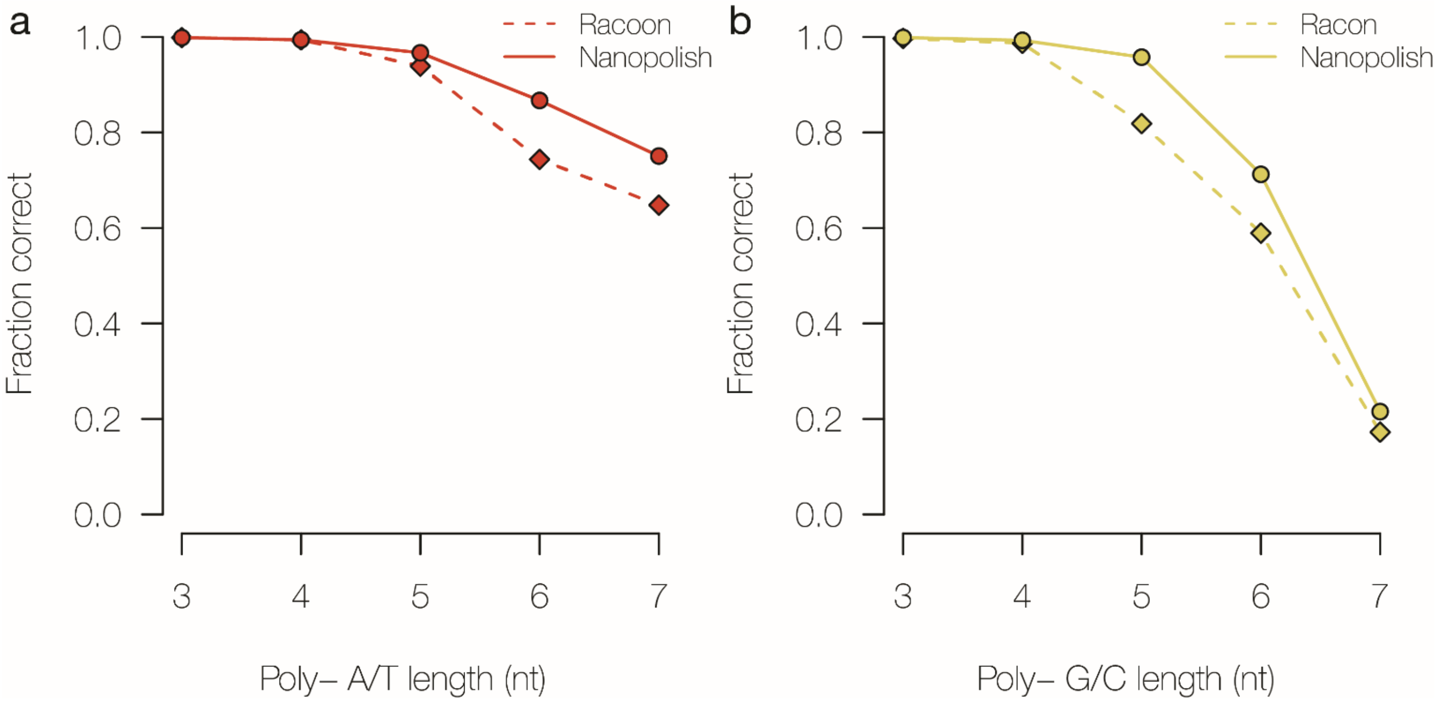

**Supplementary Figure 9. Bioinformatics pipeline for analysis of *Y. pestis* and *B. anthracis* whole genome sequencing nanopore data.** A pipeline was developed to perform rapid *de novo* genome and plasmid assembly, to detect known markers for antimicrobial resistance, and to identify variants (SNPs, insertions and deletions).

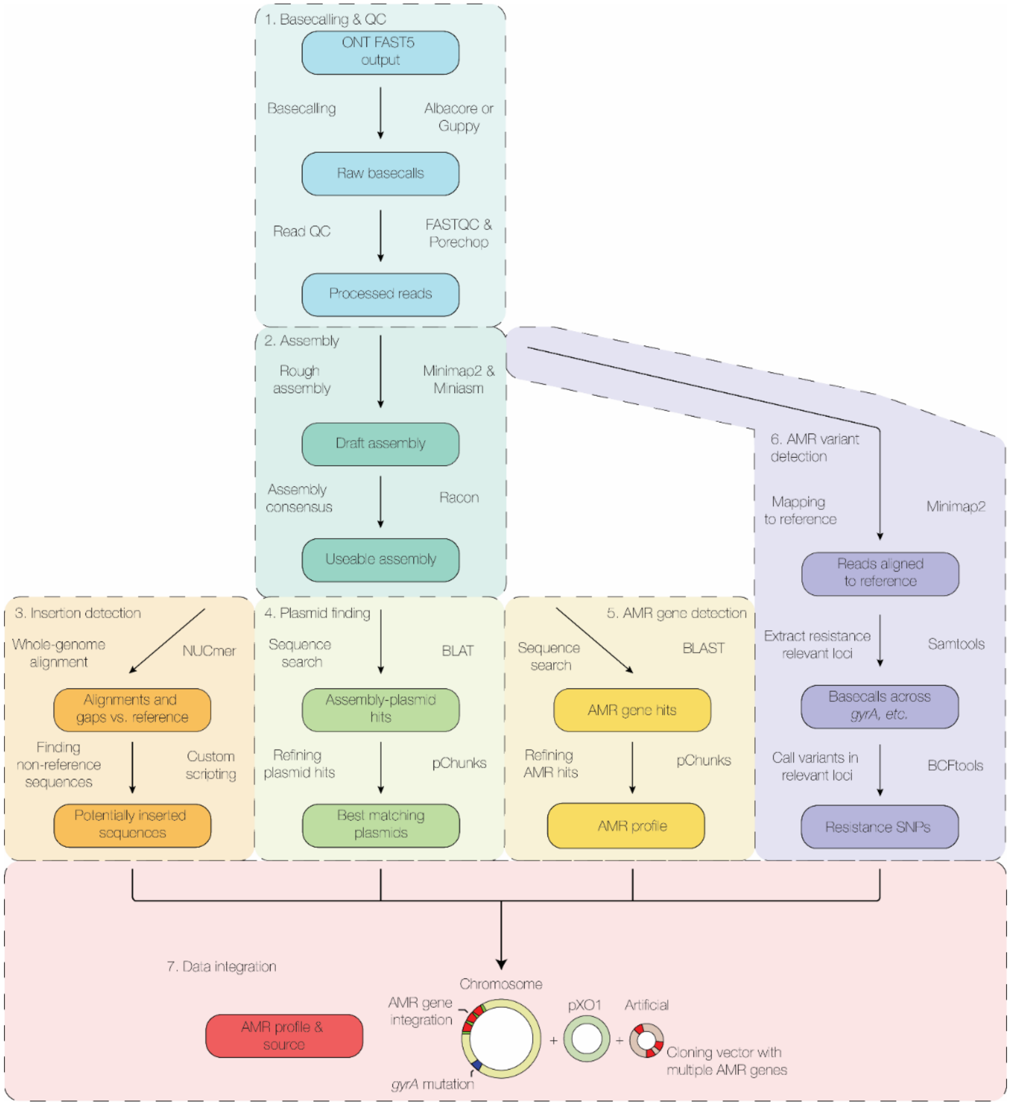
